# Hierarchical Compression Reveals Sub-Second to Day-Long Structure in Larval Zebrafish Behaviour

**DOI:** 10.1101/694471

**Authors:** Marcus Ghosh, Jason Rihel

## Abstract

Animal behaviour is dynamic, evolving over multiple timescales from milliseconds to days and even across a lifetime. To understand the mechanisms governing these dynamics, it is necessary to capture multi-timescale structure from behavioural data. Here, we develop computational tools and study the behaviour of hundreds of larval zebrafish tracked continuously across multiple 24-hour day/night cycles. We extracted millions of movements and pauses, termed bouts, and used unsupervised learning to reduce each larva’s behaviour to an alternating sequence of active and inactive bout types, termed modules. Through hierarchical compression, we identified recurrent behavioural patterns, termed motifs. Module and motif usage varied across the day/night cycle, revealing structure at sub-second to day-long timescales. We further demonstrate that module and motif analysis can uncover novel pharmacological and genetic mutant phenotypes. Overall, our work reveals the organisation of larval zebrafish behaviour at multiple timescales and provides tools to identify structure from large-scale behavioural datasets.

## Introduction

To survive, animals must coordinate patterns of action and inaction in response to their environment. These actions and inactions, which together we will define as behaviour, result from some function incorporating internal (e.g. transcriptional, hormonal or neuronal activity) and external (e.g. time of day or temperature) state. Thus, behavioural descriptions provide insight into the underlying mechanisms that control behaviour and are a necessary step in understanding these systems (Krakauer *et al.*, 2017).

Animal behaviour, however, typically has many degrees of freedom and evolves over multiple timescales from milliseconds (Wiltschko *et al.*, 2015) to days (Proekt *et al.*, 2012; Fulcher and Jones, 2017) and even across an animal’s entire lifespan (Jordan *et al.*, 2013; Stern *et al.*, 2017). As such, quantitatively describing behaviour remains both conceptually and technically challenging (Berman, 2018; Brown and de Bivort, 2018). Inspired by early ideas from ethology (Lashley, 1951; Tinbergen, 1963), one approach is to describe behaviour in terms of simple modules that are arranged into more complex motifs. Behavioural modules are often defined from postural data as stereotyped movements, such as walking in *Drosophila* (Berman *et al.*, 2014; Vogelstein *et al.*, 2014; Robie *et al.*, 2017) and mice (Wiltschko *et al.*, 2015), while behavioural motifs are defined as sequences of modules, which capture the patterns inherent to animal behaviour, such as grooming in *Drosophila* (Berman *et al.*, 2014, 2016).

Zebrafish larvae have emerged as a powerful model organism in neuroscience, owing to their genetic tractability (Howe *et al.*, 2013), translucency (Vanwalleghem *et al.*, 2018) and amenability to pharmacological screening (Rihel and Ghosh, 2015). In terms of behaviour larvae exhibit an alternating sequence of movements and pauses, termed bouts. This structure is particularly suited to modular description as individual bouts can be easily segmented and it is relatively easy to acquire many examples from even a single animal due to the high frequency of their movement (Kim *et al.*, 2017). Leveraging these advantages, recent work used unsupervised learning to uncover a locomotor repertoire of 13 swim types in larval zebrafish, including slow forward swims and faster escape swims (Marques *et al.*, 2018). However, the inactive periods between swim bouts, were not considered, despite reflecting behavioural states such as passivity in the face of adversity (Mu *et al.*, 2019) or even sleep (Prober *et al.*, 2006).

To explore an animal’s full behavioural repertoire, from fast movements to sleep it is necessary to study behaviour over long timescales. To date, however, module and motif descriptions of behaviour have been developed from videos fifteen minutes (Vogelstein *et al.*, 2014; Wiltschko *et al.*, 2015; Robie *et al.*, 2017) to two hours (Marques *et al.*, 2018) in length. Consequently, most identified behavioural structure has been on the order of milliseconds and the existence of longer-timescale structure, on the order of minutes to hours has remained largely unexplored. The development of methods to extract multi-timescale structure from long-timescale recordings would open avenues to explore questions including how behaviour varies across the day/night cycle and develops across an animal’s lifespan. Furthermore, as pharmacologically or genetically induced behavioural phenotypes can differ at different times of the day/night cycle in zebrafish larvae (Rihel *et al.*, 2010; Hoffman *et al.*, 2016), a long-timescale approach would provide valuable phenotyping information.

Currently, the limiting factor in scaling these methods is the volume of data, owing to the high-framerates and -dimensionality required to estimate animal posture. Here, we present an alternative approach in which we trade dimensionality for scale, by building a module and motif description of larval zebrafish behaviour from a one-dimensional behavioural parameter recorded over time. Specifically, we used a high-throughput behavioural set-up (Rihel *et al*., 2010) to continuously monitor the activity of hundreds of zebrafish larvae across multiple days and nights. To identify multi-timescale behavioural structure, we developed a three-step computational approach. Firstly, we used unsupervised learning to identify a set of 10 behavioural modules that describe both active and inactive bout structure. Secondly, we applied a compression algorithm (Nevill-Manning and Witten, 2000) to our module data to compile a library of almost 50,000 motifs, revealing behavioural patterns organised across sub-second to minute timescales. Finally, we used a supervised learning algorithm (Peng *et al.*, 2005) to identify motifs from the library, used at particular times of the day/night cycle. To test the ability of our approach to detect biologically relevant phenotypes, we also studied the behaviour of larvae exposed to the seizure-inducing drug, pentylenetetrazol (PTZ) (Baraban *et al.*, 2005), the sedating drug, melatonin (Zhdanova *et al.*, 2001), and hypocretin receptor (*hcrtr*) mutant larva (Yokogawa *et al.*, 2007), loss of which is associated with narcolepsy in humans (Lin *et al.*, 1999) and altered bout structure in zebrafish (Yokogawa *et al.*, 2007; Elbaz *et al.*, 2012). We found that our computational approach could readily detect both compound dose and mutant specific differences in module and motif usage, demonstrating the biological relevance of our behavioural description.

Ultimately, our work reveals the organisation of larval zebrafish behaviour at sub-second to day-long timescales and provides new computational tools to identify structure from large-scale behavioural datasets.

## Results

### Behaviour at Scale

Larval zebrafish behaviour consists of an alternating sequence of movements and pauses, termed bouts, that are organised at sub-second timescales. To capture this structure from high-throughput, long-timescale experiments, we used a 96-well plate set-up with a single larva housed in each well (Supplementary Figure 1a) and as a proxy for movement recorded the number of pixels that changed intensity within each well between successive pairs of frames, a metric we term Δ pixels. We built on previous work using this set-up (reviewed in: Barlow and Rihel, 2017; Oikonomou and Prober, 2017) by analysing Δ pixels data at 25Hz, rather than in one-minute bins. When recorded in this way, Δ pixels data is an alternating sequence of positive values representing movement magnitude and zeros representing periods of inactivity (Figure 1a, Supplementary video 1). We defined active bouts as any single or consecutive frames with non-zero Δ pixels values and described each bout using several features including the mean and standard deviation of Δ pixels values across the bout (Figure 1a). We defined inactive bouts as any single or consecutive frames with zero Δ pixels values, and described each inactive bout using its length (Figure 1a).

**Figure 1.**
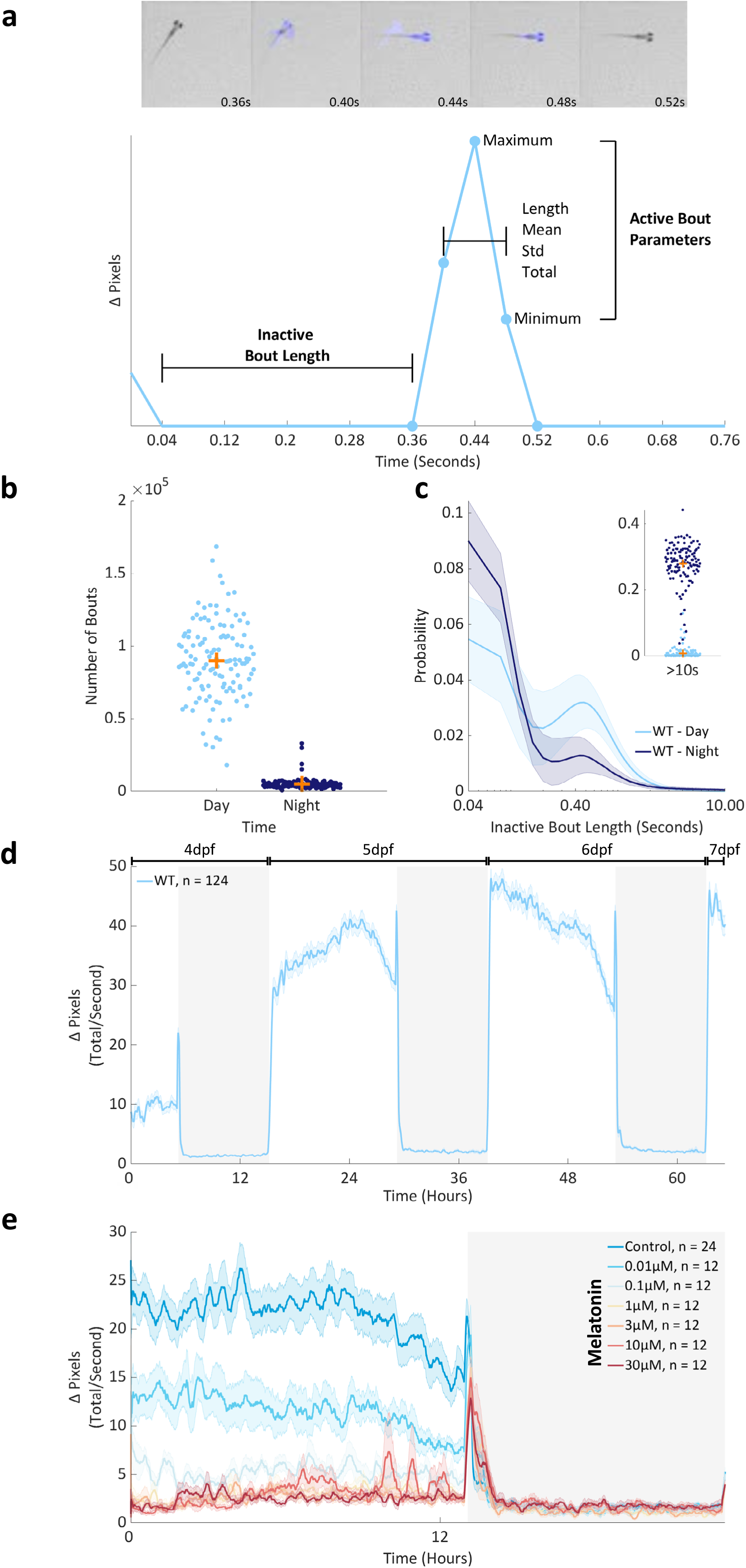
Behaviour at Scale. **a.** Top panel: five consecutive frames from an individual well of a 96-well plate as a 6dpf zebrafish larva performs a swim bout. Blue highlights pixels that change intensity between frames (Δ pixels). Lower panel: a Δ pixels time series from the larva above. Highlighted are the features that describe each active and inactive bout. **b.** The mean number of bouts recorded from individual larvae at 5 and 6dpf during the day (light blue) and the night (dark blue). Each dot is 1 of 124 wild-type larvae. The orange crosses mark the population means. **c.** The probability of observing different lengths of inactivity during the day (light blue) or the night (dark blue) at 5 and 6dpf. Each larva’s data was fit by a probability density function (pdf). Shown is a mean pdf (bold line) and standard deviation (shaded surround) with a log scale on the x-axis cropped to 10 seconds. Insert: the total probability of inactive bout lengths longer than 10 seconds, per animal. **d.** The mean activity of 124 wild-type larvae from 4-7dpf, on a 14hr/10 hr light/dark cycle. Data for each larva was summed into seconds and then smoothed with a 15-minute running average. Shown is a summed and smoothed mean Δ pixels trace (bold line) and standard error of the mean (shaded surround). **e.** Average activity across one day (white background) and night (dark background) for larvae dosed with either DMSO (control) or a range of melatonin doses immediately prior to tracking at 6dpf. Data was summed and smoothed as in d. The number of animals per condition is denoted as n=.

Using this approach, we first assessed the behaviour of wild-type larvae across a 14hr/10hr day/night cycle (Supplementary Figure 2a). During the day, wild-type larvae had many more bouts than the night (Figure 1b) and tended to use short, sub-second long inactive bouts (Figure 1c). Longer inactive bouts, on the order of seconds to minutes, were generally reserved for the night (Figure 1c). Together these differences in active and inactive bout usage resulted in a diurnal pattern of activity (Figure 1d). These results are broadly consistent with those from analysis of binned Δ pixels data (Barlow and Rihel, 2017; Oikonomou and Prober, 2017), with the addition of sub-second resolution and an increase in accuracy, as determined by intra-fish comparisons between the methods (Supplementary Figure 1b-c).

Next, we extended our approach to examine the behavioural effects of pharmacological and genetic manipulations. Melatonin, which is strongly hypnotic in zebrafish (Rihel *et al.*, 2010), dose dependently decreased larval activity (Figure 1e) by decreasing the number, magnitude, and length of active bouts and by inducing longer inactive bouts (Supplementary Figure 2b). The epileptogenic drug PTZ (Supplementary Figure 1d) altered both active and inactive bout parameters (Supplementary Figure 2c), eliciting on average longer, lower amplitude active bouts and longer inactive bouts during the day. Finally, homozygous *hcrtr*^-/-^ mutants had only subtle differences in active bout structure, with shorter mean active bout length and lower active bout total and standard deviation, compared to both wild-type *hcrtr*^+/+^ and heterozygous *hcrtr*^-/+^ siblings, which did not differ from one another by any metrics (Supplementary Figure 2d).

Collectively, these results quantitatively demonstrate the advantages of assessing Δ pixels data on a frame by frame basis and provide insight into the behaviour of wild-type zebrafish larvae across the day/night cycle as well as those subject to pharmacological or genetic manipulations.

### Module Usage Varies with Behavioural Context

Recent work has demonstrated that larval activity can be classified using unsupervised learning into 13 distinct bout types that represent different swimming movements (Marques *et al.*, 2018). A full description of larval behaviour, however, requires quantification of both the movements and pauses that they execute. Thus, we sought to determine if distinct active or inactive bout types, which we termed modules, were identifiable from our data, and if module usage depended upon behavioural context.

To address these questions, we separately clustered the active and inactive bouts (combined across experiments a total of 30,900,018 active and 30,900,418 inactive bouts) using an evidence accumulation-based clustering algorithm (see Materials & Methods). In brief, 200 Gaussian Mixture Models were built from each data set, then the results of these models were combined to generate aggregate solutions. This clustering method identified 5 active and 5 inactive modules (Figure 2a-b, Supplementary Figure 3), which we separately labelled from 1-5 from the shortest to longest mean bout length. The active modules, which formed discrete peaks in bout feature space (Supplementary Figure 3b), corresponded to different shapes of Δ pixel changes in terms of amplitude and length (Figure 2a and Supplementary Figure 4a), while the inactive modules consisted of different lengths of inactivity (Figure 2b and Supplementary Figure 4a). The shortest inactive module (module 1) had a mean length of 0.06s and ranged from a minimum of 0.04s (our sampling limit) to a maximum of 0.12s. In contrast, the longest inactive module (module 5) had a mean length of 96s and covered a huge range of values from a minimum of 20s to a maximum of 8.8hours.

**Figure 2.**
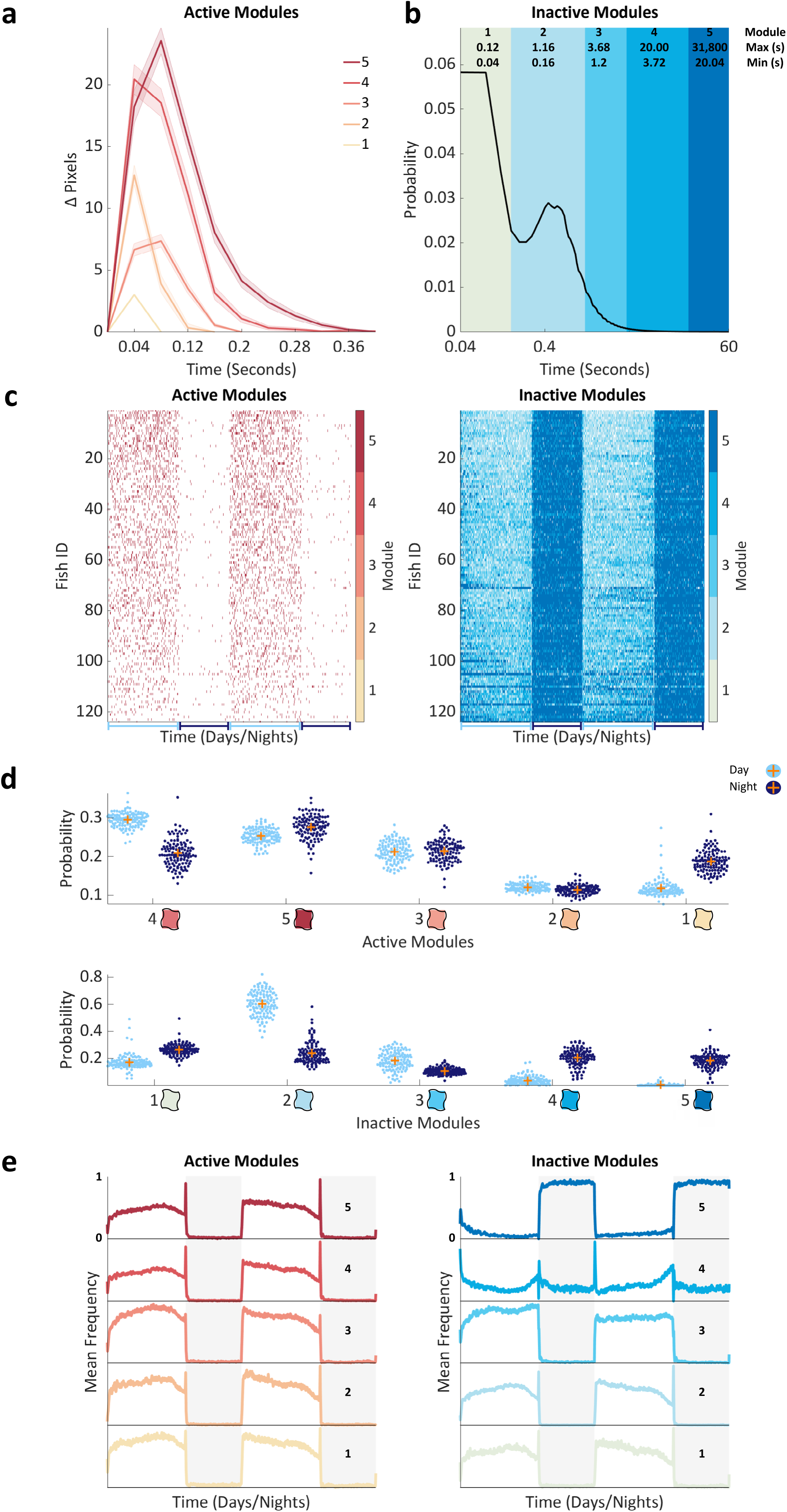
Unsupervised Learning Identifies Contextual Behavioural Modules. **a.** Average Δ pixels changes for each active module. Shown is the mean (bold line) and standard error of the mean (shaded surround) of 100 bouts randomly sampled from each module from one representative larva. Modules are numbered and coloured by average module length across all animals, from shortest (1) to longest (5). **b.** A probability density curve showing the distribution of inactive bout lengths in seconds, on a log x-axis cropped to 60s. Modules are numbered and coloured from shortest (1) to longest (5) mean length (see legend for each modules minimum and maximum bout length). **c.** Matrices showing the active (left) or inactive (right) module assignment of every frame (x-axis) for each of 124 wild-type larvae (y-axis) across the 14-hour days (light blue underlines) and 10-hour nights (dark blue underlines) from 5-6 dpf. Larvae were sorted by total number of active modules from highest (top) to lowest (bottom). Modules are coloured according to the adjacent colormaps. **d.** Average active (upper) and inactive (lower) module probability during day (light blue) and night (dark blue) 5 and 6 of development. Each of 124 wild-type animals is shown as a dot and orange crosses mark the population means. Active modules are sorted by mean day probability from highest to lowest (left to right). Inactive modules are sorted by mean length from shortest to longest (left to right). The blobs correspond to the colour used for each module in other figures. **e.** The mean frequency of each active (left) and inactive (right) module across days 5 and 6 of development. Shown is a mean smoothed with a 15-minute running average, rescaled to 0-1. Days are shown with a white background, nights with a dark background. Modules are sorted from shortest to longest (lower to upper panels).

To examine how module usage varied across time, we represented each larvae’s behaviour as an alternating sequence of active and inactive modules (Figure 2c, Supplementary video 2). In the wild-type data, module usage varied with both time of day and development (Figure 2d). For example, the probability of observing inactive module 2, which consists of typical day pause lengths (0.16 – 1.16s), was on average 0.6 during the day and only 0.24 during the night, when inactive modules 1, 4 and 5 became more likely (Figure 2d). To reveal finer-grain temporal dynamics, we also examined each module’s mean frequency over time (Figure 2e). In general, both the active and the short inactive modules had high frequencies during the day, peaking at the light/dark transition as the larvae responded to the sudden change in illumination. In contrast, the only module with a peak in frequency at the dark-to-light transition was inactive module 4 (3.72 – 20s), which also had an increased frequency approaching the light-to-dark transition. Together these results reveal that zebrafish employ different bout types in a time of day/night dependent manner.

Next, we examined the impact of pharmacological and genetic manipulations upon bout type usage. Larvae dosed with melatonin showed a shift towards using shorter active modules and longer inactive modules (Supplemental Figure 4b). In PTZ dosed larvae, there were also shifts in active module probability. Particularly notable was the complete exclusion of active module 1 in 27 of the 28 (96.4%) PTZ dosed larvae, while control larvae used this module with 0.12 probability during the day and 0.22 during the night (Supplementary Figure 4c). These shifts likely reflect the chaotic, seizure-like swimming observed in PTZ-treated larvae (Baraban *et al.*, 2005), although no single active module clearly captured these behavioural seizures. PTZ also increased the probability of the shortest inactive (module 1) as well as the two longest inactive modules (modules 4 and 5), the latter of which are likely to correspond to the inter-ictal bouts of inactivity associated with seizures (Supplementary Figure 4c). Conversely, *hcrtr* mutants exhibited no differences in either active or inactive module probabilities compared to their wild-type siblings (Supplementary Figure 4d), demonstrating that bout type usage is similar between these mutants and wild-type animals across the day/night cycle.

Collectively, these results reveal that zebrafish behaviour in this assay can be described by 5 types of active and 5 types of inactive modules, the usage of which varies with behavioural context. Interestingly, in many contexts, both active and inactive module probabilities were shifted, suggesting that these module types may co-vary, perhaps by being arranged into recurrent sequences.

### Hierarchical Compression Reveals Structure in Zebrafish Behaviour

From a set of behavioural modules, an animal could structure their behaviour in a range of ways. At one end of this spectrum, successive modules could be organised completely randomly, such that prior modules exert no influence on future module selection. At the other end, module selection could be fully deterministic with a particular module always following another. Rather than being fixed, however, it is likely that animals adapt their behavioural structure in response to changing internal or external states. We sought to map the structure of zebrafish behaviour in different contexts by examining the presence and organisation of module sequences, which could provide insight into the mechanisms governing behaviour. To do this, we used a compression algorithm (Nevill-Manning and Witten, 2000) as Gomez-Marin and colleagues (2016) used to discover structure in *C. elegans* postural data. When applied to our dataset (Figure 3a), this algorithm iteratively identified motifs from each larva’s modular sequence and returned two outputs -- compressibility, a measure of each larva’s behavioural structure, and a library of identified recurrent module sequences, termed motifs.

**Figure 3.**
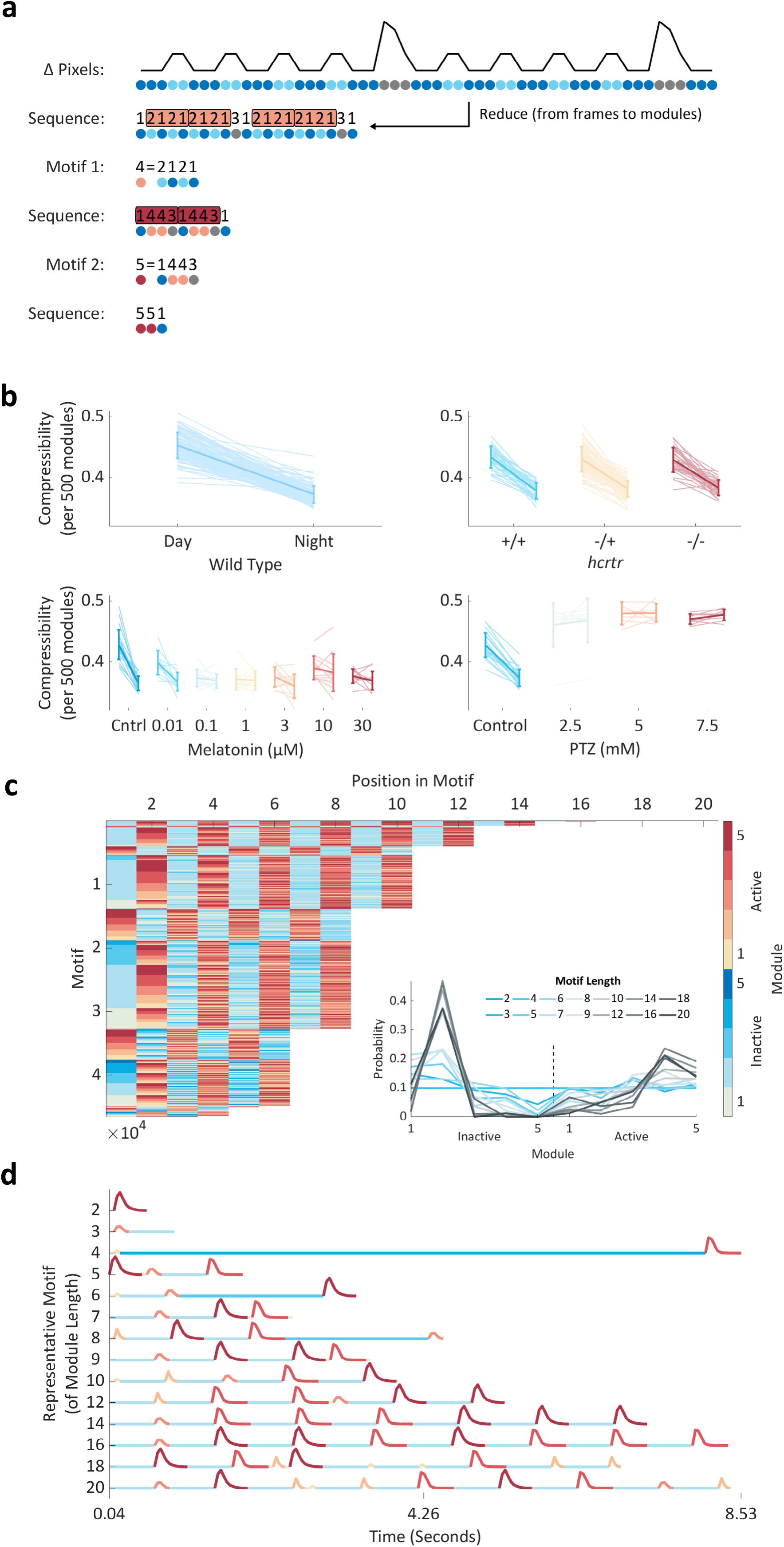
Hierarchical Compression Reveals Structure in Zebrafish Behaviour. **a.** Compression explained using fictive data. Top to bottom: from Δ pixels data (black trace) we classified both active and inactive behaviours into modules (coloured circles). From modular behavioural sequences, we identified motifs (sequences of modules) using a compression algorithm. Compression iteratively identifies motifs (shown as boxes) by replacing them with new symbols until no more motifs can be identified and the sequence is maximally compressed. **b.** Each panel shows how compressibility, calculated from 500 module blocks, varies in different behavioural contexts. Each pale line shows an individual fish’s mean compressibility during the day and the night. The darker overlay shows a population day and night mean ± standard deviation. In the wild-type data, compressibility is higher during the day than the night (p < 10^-158^) and increases from day/night 5 to 6 (p < 10^-4^), findings consistent across triplicate experiments. Melatonin decreases (p < 10^-10^), while PTZ increases compressibility (p < 10^-8^). There is no effect of *hcrtr* genotype on compressibility. Statistics are two or four-way ANOVA. **c.** All 46,554 unique motifs (y-axis) identified by compressing data from all animals. Each motif’s module sequence is shown, with the modules coloured according to the colormap on the right. Motifs are sorted by length and then sequentially by module. Motifs range in length from 2-20 modules long. Insert: for each motif length, the probability of observing each inactive or active module. **d.** Each motif in the library consists of an alternating sequence of Δ pixels changes and pauses (active and inactive modules). A representative motif of each module length is shown with each module coloured according to the colormap in c. Representative motifs were chosen by determining every motif’s distribution of modules and then for each observed module length, selecting the motif closest to the average module distribution (see c, insert).

To quantify the structure of zebrafish behaviour, we first compressed every animal’s full modular sequence, which in wild type animals were on average 236,636 modules long across 70 hours. To determine if the resultant compression values indicated more structure than would be expected based on either the distribution or the transition structure of the active-to-inactive modules, we compared each larva’s compressibility to that of 10 sets of paired shuffled data. All wild-type larvae were more compressive than their paired shuffled data, demonstrating that their behaviour is more structured than expected from modular probabilities alone (Supplementary Figure 5a). Compressibility, however, varies non-linearly with input sequence length, as longer sequences will be more likely to contain motifs (Supplementary Figure 5b). Thus, to enable comparisons between samples with different numbers of modules, we compressed non-overlapping 500 module blocks of sequence per larva. This approach revealed that compressibility was higher during the day than the night (Figure 3b) and increased with developmental age. To determine if these differences were primarily due to the presence of behavioural motifs or instead were a consequence of differences in module distribution, we also compared the difference in compressibility (Δ compressibility) between each animal’s real and shuffled data. This approach revealed that the compressibility difference between the day and the night is predominantly due to differences in module selection (Supplementary Figure 5d). To reveal finer-grain temporal changes in compressibility, we plotted Δ compressibility across time (Supplementary Figure 5e). This approach revealed peaks at the light-to-dark transitions in the evenings, consistent with this stimulus eliciting stereotyped behavioural sequences (Burgess and Granato, 2007; Emran *et al.*, 2010).

Next, we used compressibility to assess how our pharmacological and genetic manipulations altered the structure of larval behaviour. We found that melatonin decreased day compressibility to night-time levels (Figure 3b). In contrast, PTZ increased compressibility to a constant day/night value (Figure 3b). PTZ, however, reduced Δ compressibility (Supplementary Figure 5d), indicating that changes in module distribution, rather than motif usage, are the dominant driver of PTZ-induced behavioural changes. Importantly, these drug-induced changes in compressibility do not simply reflect overall activity levels. For example, PTZ exposed larvae are less active than controls during the day and more active during the night (Supplementary Figure 1d) but have consistently higher compressibility (Figure 3b). Finally, in *hcrtr* mutants we found no differences in either compressibility or Δ compressibility, suggesting that *hcrtr* mutant behaviour is structured similarly to wild-type animals (Figure 3b).

To gain insight into the behavioural sequence’s larvae deploy, we then studied the motifs identified by the compression algorithm. Compression of the real modular sequences identified a mean of 1901 motifs per animal (Supplementary Figure 5c). Interestingly, compression of the real data almost always identified slightly fewer motifs than the shuffled data (Supplementary Figure 5c). This suggests that the motifs identified from the real data were used more frequently than those in the shuffled data and therefore likely reflect enriched behavioural sequences. Merging the motifs identified across all animals generated a library of 46,554 unique behavioural motifs (Figure 3c). In terms of raw Δ pixels data, each motif represented an approximately repeated pattern of movements and pauses of varying length (Figure 3d). Motifs in the library ranged from 2-20 modules long with a median length of 8 modules and spanned timescales from approximately 0.1s-11.3 minutes with a median length of 3.84s. Motifs of different module lengths used distinct sub-sets of modules (Figure 3c). For example, motifs comprised of longer module sequences had a lower probability of using long inactive modules. Together, these results reveal the varied timescales at which zebrafish larvae organise their behaviour and suggest the presence of structure governing the arrangement of modules into motifs.

### Behavioural Motif Usage is Time Dependent

The large number of motifs in our library led us to hypothesise that each may be used in specific behavioural contexts. To test this hypothesis, we counted the number of times each larva used each motif within each time frame (e.g. day or night) and then normalised these counts by calculating whether each motif was observed more or less frequently than in the paired shuffled data, a metric we termed enrichment/constraint. Overall, we found that enrichment/constraint scores from our real data were more prone to extreme positive (enriched) and negative (constrained) values than the shuffled data (Figure 4a), suggesting that a minority of behavioural motifs were used more or less frequently than would be expected by chance.

**Figure 4.**
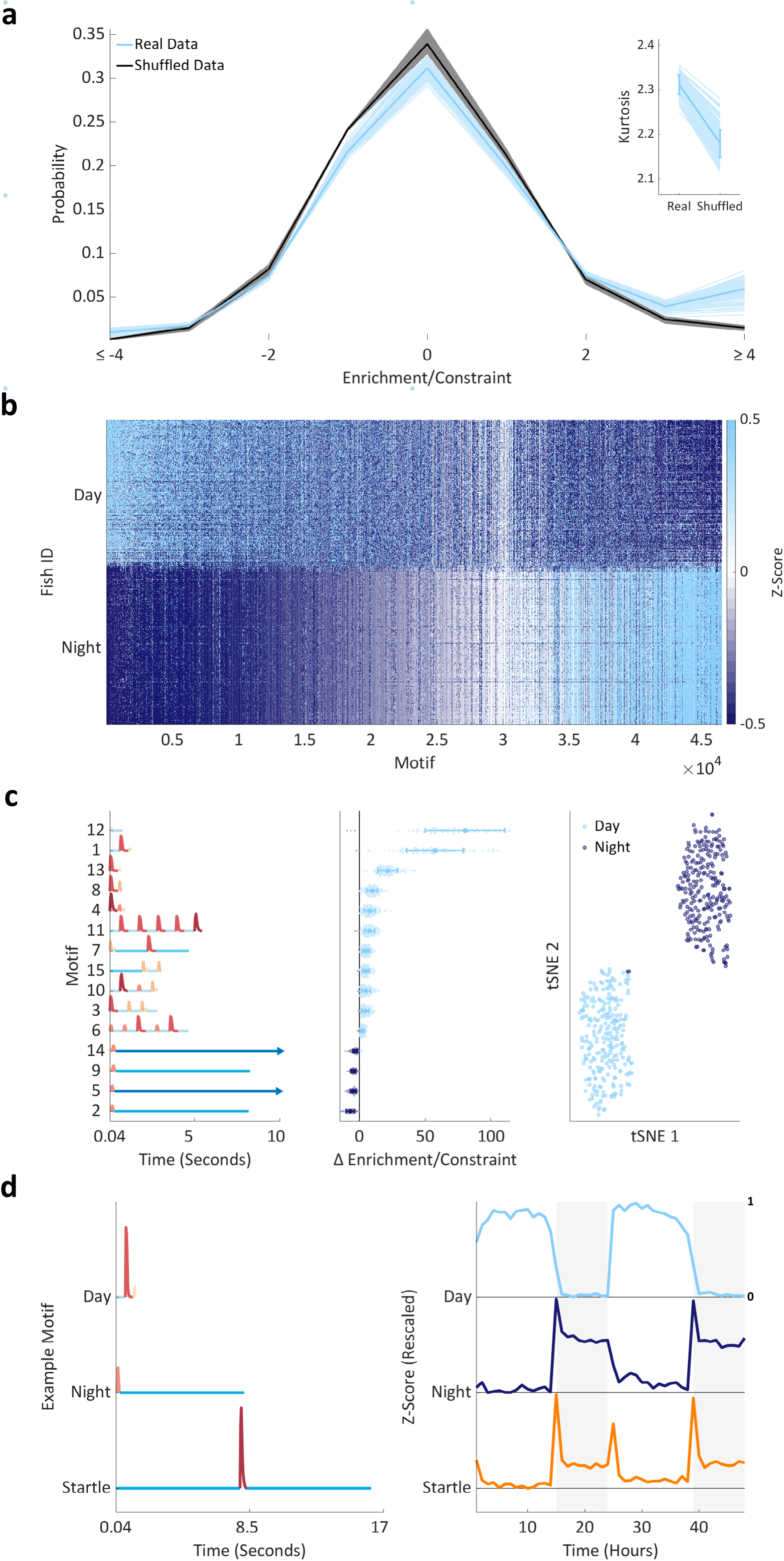
Supervised Learning Identifies Contextual Behavioural Motifs. **a.** Probability density functions (pdfs) showing the probability of observing motifs at different enrichment/constraint scores rounded to whole numbers and summed at values above or below ± 4 for ease of visualisation. Each wild type animal is depicted by a single pale blue (real data) and 10 black (shuffled data) lines; overlaid in bold are mean pdfs. The insert shows that the kurtosis of the real data is higher than the shuffled data (p < 10^-271^; two-way ANOVA, real vs shuffled data, no significant interaction with experimental repeat factor). Each larva is shown as a pale line; overlaid is a population mean and standard deviation. **b.** Enrichment/constraint scores for all 46,554 motifs (x-axis) for each fish during day/night 5 and 6 of development (y-axis). To emphasise structure, motifs are sorted in both axes, first by their average day night difference (from day to night enriched left to right), then separately day and night by larva. Finally, each motif’s enrichment/constraint score is Z-scored to aid visualisation. **c.** Left: the 15 day/night mRMR motifs module sequences are shown numbered by the order in which they were selected by the algorithm. Motifs are sorted by day minus night enrichment/constraint score (middle). The long pauses at the end of motifs 5 and 14 are cropped at 10s (arrows). Middle: for each selected motif (y-axis), ordered as in the left panel, each wild-type animal’s (124 in total) day minus night enrichment/constraint score (x-axis) is shown as a dot. Values above zero are coloured light blue; below zero are dark blue. Overlaid is a population mean and standard deviation per motif. Right: a tSNE embedding of the 15-dimensional motif data (middle) into a 2-dimensional space. Each circle represents a single day (light blue) or night (dark blue) sample. **d.** Representative motif temporal dynamics; shown are motifs 1 (day) and 2 (night) from c, as well as a startle-like motif. Left: each motif’s module sequence. Right: each motif’s mean enrichment/constraint score each hour, rescaled to 0-1.

To test if these extremes occurred in particular contexts, we first compared motif usage between the day and the night in wild-type larvae by generating a matrix of enrichment/constraint scores (Figure 4b). To distil the most salient motifs from this and other contextual matrices, we used a three-step approach. Firstly, we used the minimal-redundancy-maximal-relevance criterion (mRMR) algorithm (Peng *et al.*, 2005) to rank the motifs from most to least salient. Secondly, we trained linear discriminant analysis classifiers using 10-fold cross validation as we iteratively increased the number of input motifs from most to least salient (e.g. motif 1, motif 1 & 2, motif 1 – n). Finally, we selected the subset of motifs which achieved the lowest classification error between groups in each context. To determine how accurately these motif subsets could distinguish between behavioural contexts, we compared each classifier’s performance to that of a majority class classifier, which performed as well as the ratio of samples between the two contexts. For example, in the day vs. night classification, a majority class classifier would have an error rate of 50% (± standard error of proportion), as each larva contributes an equal number of days and nights to the enrichment/constraint matrix. Additionally, to demonstrate the salience of the motifs selected by the mRMR algorithm, we compared each classifiers performance to a set of 10 classifiers built using the same number of motifs, though randomly selected. For example, for a classifier which achieved its minimal classification error using 50 motifs, we randomly selected 50 motifs from the library and built a classifier. For each comparison we repeated this process 10 times.

Applying this algorithm to wild-type data revealed changes in motif usage across multiple timescales (Supplementary Figure 6b). We found that only 15 motifs were required to classify day- and night-specific behaviour with only a 0.2% (±0.63% Std) classification error, compared to a majority class classifier with 50% error and random 15-motif subsets with a mean error of 9.25% (Figure 4c, Supplementary Table 1). The day enriched motifs consisted of high amplitude movements interspersed with short pauses, while the night enriched motifs contained low amplitude movements and long pauses (Figure 4c). Next, we examined how motif usage changed over development by comparing consecutive days and nights (5-6dpf). In both day 5 vs. day 6 and night 5 vs. night 6 comparisons, the classifiers achieved roughly 20% error using 93 and 85 motifs, respectively (Supplementary Table 1). Thus, motif usage shifted over just 24 hours of development, though these changes were far less prominent than those between the day and night. To study whether motif usage varied at finer timescales, we first divided the day into morning/evening and the night into early/late periods. In each case the mRMR algorithm performed better than the control classifiers (morning/evening: 33%, early/late night: 36%) though the relatively high classification errors suggest that motif selection did not vary strongly across each day or night (Supplementary Table 1). Consistent with this conclusion, classifiers attempting to delineate each hour from every other mostly failed to outperform their control classifiers (Supplementary Table 1). The two notable exceptions were the hour following each lighting transition, where this approach identified motifs with startle-like patterns (Figure 4d) and achieved good classification performance (Supplementary Table 1). Together these results demonstrate that motif usage varied between the day and the night, but aside from the lighting transitions, was relatively consistent within these periods.

### Dose-Dependent and Dose-Specific Behavioural Motifs

Finally, we hypothesised that behavioural motif usage would vary dose-dependently across concentrations of melatonin and PTZ, providing insight into the mechanisms by which these compounds exert their behavioural effects. Motif dose-dependency would suggest a continuously modulated underlying process, which might arise if the fraction of bound receptors relates to neuronal activity modulation. Alternatively, motifs enriched at only specific doses, would suggest discrete effects upon neuronal circuitry, for example the binding of low affinity receptors.

Applying the mRMR algorithm to our pharmacological data revealed both dose-dependent and dose-specific modulation of motif usage. We found that each melatonin dose could be separated from the others using 40 to 250 motifs with only 0-2.78% classification error (Figure 5a, Supplementary Table 2). Focussing on just the best motif for each comparison, we observed both dose-dependency as well as dose-specificity. For example, comparing controls to all melatonin-dosed larvae identified a dose-dependent motif that consisted of large magnitude movements and short pauses, whose enrichment/constraint score decreased with increasing melatonin concentration (Figure 5a). Conversely, the best 10µM motif, two long pauses broken by a small active bout sequence, showed dose-specificity being enriched at only 3µM and 10µM doses (Figure 5a). When applied to the PTZ data, our approach performed even more accurately, achieving perfect classification (0% error) between all conditions (Figure 5b and Appendix Table 2). Furthermore, in PTZ-dosed larvae we observed enrichment for motifs highly constrained in wild-type larvae, highlighting the usage of motifs beyond the normal wild-type repertoire, such as those corresponding to behavioural seizures (Figure 5b).

**Figure 5.**
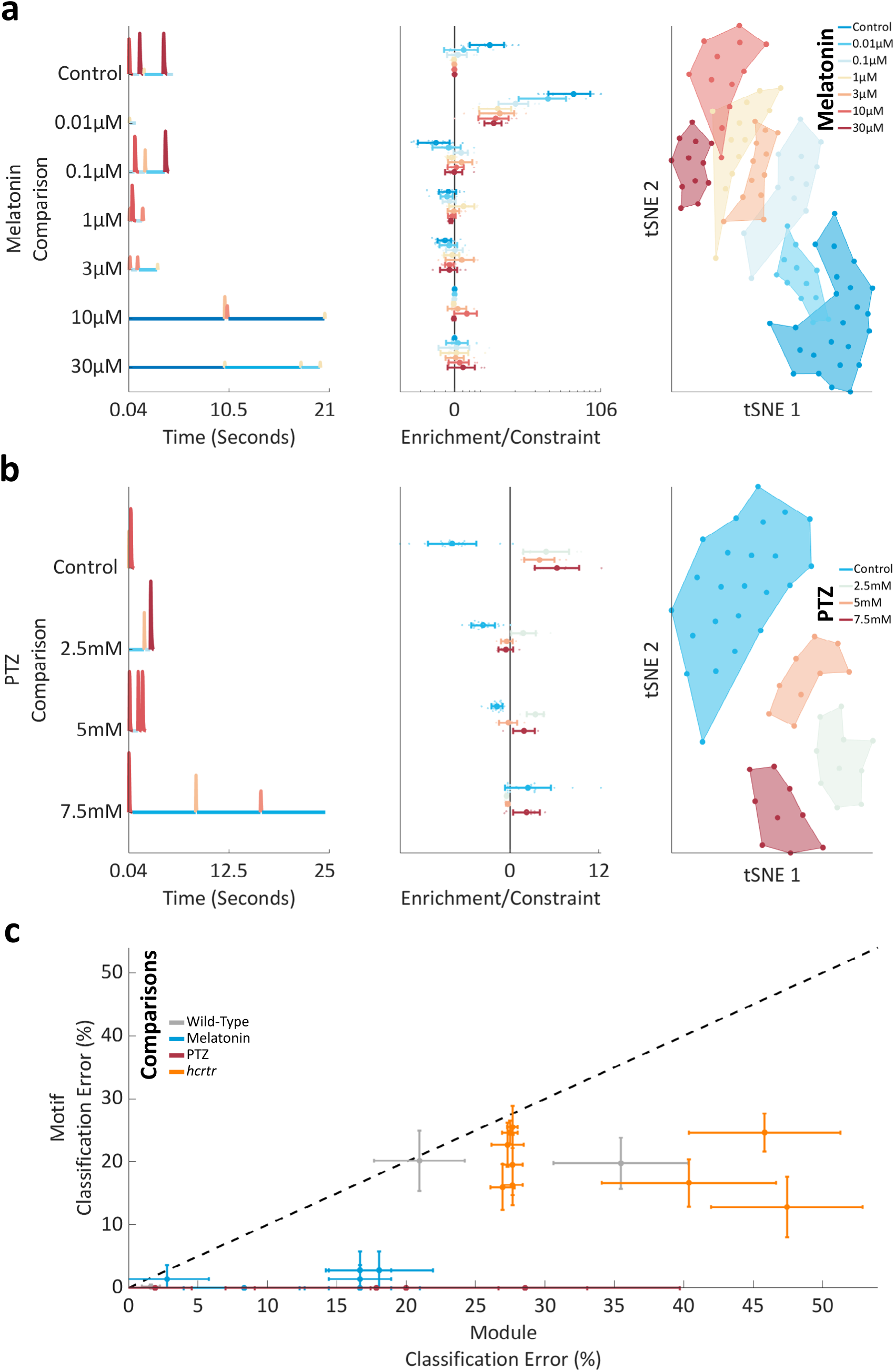
Pharmacological Behavioural Motifs. **a.** Left: module sequences for the single best motif for each melatonin comparison. Modules are coloured as elsewhere. Middle: for each dose’s single best motif, see left panel y-axis for dose, enrichment/constraint scores are shown for every dose on a log x-axis. Each animal is shown as a dot, with a mean ± std overlaid per dose. Right: a 2-dimensional tSNE embedding from a space of 912 unique motifs. Each animal is shown as a single dot underlaid by a shaded boundary encompassing all animals in each condition. **b.** Left: module sequences for the single best motif for each PTZ comparison. To highlight a seizure specific motif, the control motif and corresponding enrichment/constraint score shown is mRMR motif 2, not 1, for this comparison. Modules are coloured as elsewhere. Middle: for each dose’s single best motif, enrichment/constraint scores are shown for every dose on a linear x-axis. Each animal is shown as a dot, with a mean and standard deviation overlaid per dose. Right: a 2-dimensional tSNE embedding from a space of 338 unique motifs. Each animal is shown as a single dot underlaid by a shaded boundary encompassing all animals in each condition. **c.** Each classifier’s classification error (%) is shown in terms of modules (x-axis) and motifs (y-axis). Data is shown as mean and standard deviation from 10-fold cross validation. Classifiers are coloured by experimental dataset (see Legend). For reference, y = x is shown as a broken black line. Data below this line demonstrates superior performance of the motif classifiers.

Next, we tested whether our motif subset approach could detect *hcrtr* mutant phenotypes that were not easily captured by other methods. For example, based upon human and rodent literature, where loss of hypocretin is associated with narcolepsy (Lin *et al.*, 1999) and prior zebrafish literature (Elbaz *et al.*, 2012), we expected abnormal transitions between active and inactive bouts. We found reasonable performance when discriminating between *hcrtr*^+/+^ and *hcrtr*^-/-^ during both the day (16.7 ± 7.5% error with 195 motifs) and night (12.8 ± 9.6% error with 53 motifs) but weaker performance when distinguishing between *hcrtr*^+/+^ and *hcrtr*^-/+^, as expected for a haplosufficient gene (Supplementary Figure 6c and Supplementary Table 2). Thus, homozygous loss of *hcrtr* impacts motif usage enough to allow for successful classification of *hcrtr*^-/-^ mutants, though no single *hcrtr*^-/-^ motifs with large differences in enrichment/constraint scores compared to wild type siblings were particularly evident.

Collectively, these results demonstrate that behavioural motifs are used context dependently and reveal how motif subsets can parse subtle differences in motif usage between behavioural contexts. However, does motif analysis provide additional discriminatory power over module selection, which also varies between behavioural contexts? To assess this, we compared the performance of each motif classifier to paired module classifiers built from matrices of module probabilities. All of the motif classifiers achieved better performance than their module pairs (Figure 5c), demonstrating both the phenotyping value of the motifs and their importance in the structure of larval behaviour.

## Discussion

Here, we developed and applied computational tools to describe high-throughput, long-timescale behavioural data in terms of behavioural units (modules), and sequences of modules (motifs) organised across sub-second to day-long timescales.

### Low-Dimensional Representations of Behaviour

Low dimensional representations of behaviour, such as the Δ pixels metric employed here, result in a loss of information, for example direction of movement or posture. Such metrics do however facilitate screening approaches and/or long-timescale tracking and in these contexts have provided biological insight into the molecular targets of small molecules (Rihel *et al.*, 2010) and genetics of ageing (Churgin *et al.*, 2017). Our work builds on previous long-timescale studies of behaviour by assessing sub-second resolution Δ pixels data across multiple days and nights. This improved resolution enabled the segmentation and parameterisation of individual active and inactive bouts from our data, revealing how larvae adapt their behaviour across the day/night cycle and how behaviour is impacted by small molecules.

Future work should aim to extend our assay by recording more detailed behavioural measures. Indeed, a recent study using centroid tracking in 96 well plates revealed that larvae show a day/night location preference within the well, and furthermore uncovered a mutant with a difference in this metric (Thyme *et al.*, 2019), demonstrating that even within the confined space of a 96-well plate, location is an informative metric to record. It is likely that even more detailed behavioural measures, like eye and tail angles, will yield additional insights, for example enabling the exploration of rapid-eye-movement sleep in zebrafish larvae (Shein-Idelson *et al.*, 2016). Such metrics could be extracted by skeletonization or even through the use of an autoencoder applied to the raw video frames from each well (Johnson *et al.*, 2016).

### Modular Descriptions of Behaviour

A key idea in ethology is that behaviour consists of stereotyped modules arranged into motifs (Lashley, 1951; Tinbergen, 1963). While early studies described behaviour in this manner through manual observations (Richard and Dawkins, 1976), recent advances in machine vision and learning have automated these processes (Todd *et al.*, 2017). For example, in zebrafish larvae, recent work used unsupervised learning to uncover a locomotor repertoire of 13 swim types including slow forward swims and faster escape swims (Marques *et al.*, 2018), although inactive bouts were not considered. From our dataset, we identified 5 active and 5 inactive modules, which respectively describe swim bouts of different amplitudes (Figure 2a) and periods of inactivity of varied length (Figure 2b). Interestingly, all modules were used with reasonably high and similar probability by all wild-type animals (Figure 2d), demonstrating that these modules represent a set of common larval behaviours. Furthermore, the temporal (Figure 2e) and pharmacological (Supplementary Figure 4b-c) shifts in these probabilities illustrates that module usage can be flexibly re-organised depending upon behavioural context (Wiltschko *et al.*, 2015).

To discretize our bouts into modules, we first extracted hand-engineered features from each bout (Figure 1a) and then applied an evidence accumulation based clustering algorithm (Fred and Jain, 2002, 2005). While our results demonstrate the relevance and utility of these modules in describing larval behaviour, it is possible that our approach missed rare bout types. Consequently, future work should build upon our bout classification by exploring the benefits of including additional features, the use of alternative clustering algorithms and our assumption of stereotypy, i.e. that all bouts can be fit into a module (Berman, 2018). An alternative direction would be to produce a mapping between our active modules and those identified from analysis of larval posture (Marques *et al.*, 2018). Bridging this gap could facilitate behavioural screening approaches, for example by using data from our set-up to prioritise pharmacological compounds or mutants for postural analysis.

### Quantifying Structure in Behaviour

In some contexts, it is beneficial for animals to execute coordinated patterns of behaviour. For example, to efficiently search an environment zebrafish larvae will execute organised sequences of left and right turns (Dunn *et al.*, 2016). In other contexts, more random behaviour will be advantageous, such as when escaping from a predator (Maye *et al.*, 2007). Quantifying structure in behaviour thus provides insight into the overarching strategy being employed in particular contexts. Alterations in behavioural structure can also manifest clinically, for example in Autism Spectrum Disorder, a defining feature of which is increased behavioural stereotypy (American Psychiatric Association, 2013). Consequently, compression would be a relevant and likely informative metric to record in animal models or even human cases for such conditions.

To quantify structure in larval zebrafish behaviour in different contexts, we inputted each larva’s modular sequence to a compression algorithm. We found that wild-type behaviour was more compressive during the day than the night (Figure 3b). This echoes recent work in *Drosophila* that revealed higher temporal predictability during the day than the night as well as in females (Fulcher and Jones, 2017). A likely explanation for these findings comes from work in *C. elegans* (Gomez-Marin *et al.*, 2016) that demonstrated that animals who transition slowly between modules, as both zebrafish (Figure 1b) and *Drosophila* do at night (Geissmann *et al.*, 2019), tend to be less compressive. This may suggest that the underlying mechanisms controlling longer-timescale behaviours are less precise than those controlling fast behavioural sequences.

For future efforts applying compression to behavioural data, there are two avenues left to explore — what compression heuristic to use and how to compress data from multiple animals. Following the work of Gomez-Marin and colleagues (2016), we defined the best motif at any iteration as the most compressive, which represents a balance between the motif’s length and frequency. While this metric generally leads to the best compression (Nevill-Manning and Witten, 2000), alternative measures, such as frequency or length may capture other aspects of behaviour. The second avenue relates to comparisons between animals. Here, each animal was compressed individually, identifying motifs, which were later grouped into a common library. Whilst computationally tractable, this approach prevents certain comparisons across animals, for example identifying the most compressive motif across all larvae. This issue could be solved by compressing a single sequence containing all of the animal’s modular sequences joined end to end, with spacers to prevent inter-animal motifs. Compressing this long sequence would, however, be computationally demanding.

Compressing and merging the identified motifs across all animals generated a library of 46,554 unique motifs (Figure 3c), each of which described an alternating sequence of movements and pauses (Figure 3d). Motifs ranged from 0.1s to 11.3 minutes in length, revealing the range of timescales at which larval behaviour is organised. We cannot, however, rule out the existence of longer timescale motifs in larval behaviour as computational demands limited our search to motifs 10 modules long (though the algorithm’s hierarchical approach enabled the identification of motifs up to 20 modules long). Thus, future work should aim to extend our approach to explore the full range of timescales at which larval behaviour is organised by systematically varying this parameter.

### Contextual Behavioural Motifs

Finally, by distilling salient subsets of motifs from our library, we demonstrated that motif usage was context dependent and highlighted the discriminatory power of motif subsets, which were capable of distinguishing between day/night behaviour and even between small changes in compound dose. Comparing motif usage across the day/night cycle identified a set of highly night specific motifs (Figure 4c), which may represent sleep behaviours. One way in which future studies could address this possibility would be to deprive larvae of these motifs throughout the night, for example by using a closed-loop paradigm (Geissmann *et al.*, 2019), and observing the impact on larval behaviour the following day. In relation to the PTZ data, comparing seizure motifs across epileptogenic compounds and mutants with spontaneous seizures could suggest clues as to their underlying mechanism (Kokel *et al.*, 2010; Rihel *et al.*, 2010). For example, seizures with similar motif usage patterns may originate in the same brain area or impact awareness in the same manner. This hypothesis could be tested by generating whole-brain activity maps (Randlett *et al.*, 2015) across conditions, with the aim of identifying common and unique neuronal correlates.

Given the amenability of larval zebrafish to high-throughput behavioural screening (Rihel and Ghosh, 2015) future work should leverage our approach to large-scale genetic (Thyme *et al.*, 2019) or pharmacological datasets (Rihel *et al.*, 2010). Individually, these datasets would provide information on the genetic and molecular basis of behaviour across multiple timescales, encompassing processes from sleep to ageing. In combination, by identifying mutant and drug-induced phenotypes that cancel each other out (Lamb *et al.*, 2006; Hoffman *et al.*, 2016), these datasets could be used to identify phenotypic suppressors in genetic disease models, an outcome with potential clinical relevance.

## Materials and Methods

### Animal Husbandry

Adult zebrafish were reared by UCL Fish Facility on a 14hr/10hr light/dark cycle (lights on: 09:00 a.m. to 23:00 p.m.). To obtain embryos, pairs of adult males and females were isolated overnight with a divider that was removed at 09:00 a.m. the following morning. After a few hours, fertile embryos were collected and sorted under a bright-field microscope into groups of 50 embryos per 10 cm petri dish filled with fresh fish water (0.3g/L Instant Ocean). Plates were kept in an incubator at 28.5°C on a 14hr/10hr light/dark cycle. Using a Pasteur pipet under a bright-field microscope, debris was removed from the plates and the fish water replaced each day. All work was in accordance with the UK Animal Experimental Procedures Act (1986) under Home Office Project Licence 70/7612 awarded to JR.

### Behavioural Setup

For all behavioural experiments a Pasteur pipet was used to transfer single zebrafish larvae (aged 4-5 days post fertilisation) into the individual wells of a clear 96-square well plate (7701-1651; Whatman, New Jersey, USA); then each well was filled with 650µl of fish water. For experiments longer than 24 hours, larvae were plated at 4 days post fertilisation (dpf) and tracking was started the same day. For the duration of these experiments, evaporated fish water was replaced each morning between 09:00-09:30 a.m. For the wild-type experiments, each plate was covered with a plastic lid (4311971; Applied Biosystems, Massachusetts, USA) to prevent evaporation and to negate the need to replenish the fish water. For the 24-hour small molecule experiments (melatonin and PTZ), larvae were plated at 5dpf and the plates were left overnight in a 28.5°C 14hr/10hr light/dark incubator. The following morning each plate was transferred to a behaviour setup where larvae were dosed, between 09:00 and 10:00 a.m., immediately after which behavioural recordings were started and run for 24 hours.

To record each animal’s behaviour, each plate was placed into a Zebrabox (ViewPoint Life Sciences, Civrieux, France) running quantization mode with the following settings: detection sensitivity -- 15, burst -- 50 and freezing -- 4. All experiments were conducted on a 14hr/10hr light/dark cycle (lights on at 09:00 a.m. to 23:00 p.m.) with constant infrared illumination. All experiments were recorded at 25Hz. Larvae were tracked continuously for 24-73 hours, after which all larvae unresponsive to touch with a 10µl pipette tip were presumed sick or dead and excluded from subsequent analysis. Following this, larvae were euthanised with an overdose of 2-Phenoxyethanol (Acros Organics, New Jersey, USA).

### Fish Lines

The term “wild-type” refers to the AB x TUP LF zebrafish strain. This line was used for the wild-type experiments, as well as the melatonin and PTZ dose response curves. *hcrtr* (ZFIN ID: hu2098 (Yokogawa *et al.*, 2007). Identified from an ethylnitrosourea-mutagenized screen. UCL Line 2114.) experiments were carried out on embryos collected from heterozygous in-crosses, with larvae genotyped using KASP primers (LGC Genomics, Hoddesdon, UK) post-tracking. KASP results were validated by comparison to PCR-based genotyping of samples from each KASP classified genotype.

### *hcrtr* Genotyping

#### DNA Extraction

Following each *hcrtr* experiment each larva was euthanised in its well (as above) and DNA was extracted using HotSHOT DNA preparation (Truett *et al.*, 2000). Larval samples were transferred to the individual wells of a 96-well PCR plate. Excess liquid was pipetted from each well before applying 50µl of 1x base solution (1.25M KOH, 10mM EDTA in water). Plates were heat sealed and incubated at 95°C for 30 minutes then cooled to room temperature before the addition of 50µl of 1x neutralisation solution (2M Tris-HCL in water).

#### PCR

The following reaction mixture per sample was prepared on ice in a 96-well PCR plate: 18.3µl PCR mix (2mM MgCl_2_, 14mM pH 8.4 Tris-HCl, 68mM KCl, 0.14% Gelatin in water, autoclaved for 20 minutes, cooled to room temperature, chilled on ice, then we added: 1.8% 100mg/ml BSA and 0.14% 100mM d [A, C, G, T] TP), 0.5µl of forward and reverse primers (20 µM), 5.5µl water, 0.2µl of Taq polymerase and 3.0µl of DNA. Next, each plate was heat sealed and placed into a thermocycler, set with the following program: 95°C -- 5 minutes, 44 cycles: 95°C -- 30 seconds, 57°C -- 30 seconds and 72°C -- 45 seconds, then 72°C -- 10 minutes and 10°C until collection. Finally, samples were mixed with 6x loading buffer (Colourless buffer: Ficoll-400 - 12.5g, Tris-HCl (1M, pH 7.4) – 5ml, EDTA (0.5M) – 10mL, to 50ml in pure water; heated to 65°C to dissolve, per 10ml of colourless buffer 25mg of both xylene cyanol and orange G were added, then diluted to 6x) and run on agarose gels (1-2%) with 4% GelRed (Biotium, California, USA).

*hcrtr* Forward Primer: 5’ CCACCCGCTAAAATTCAAAAGCACTGCTAAC 3’

*hcrtr* Reverse Primer: 5’ CATCACAGACGGTGAACAGG 3’

PCR Information: PCR products were digested with Ddel at 37°C to produce a 170bp band in the wild type animals and in *hcrtr* mutants 140 and 30bp bands.

#### KASP

KASP genotyping was carried out in white, low profile PCR plates on ice with six wells allocated 50:50 for positive and negative controls. The following reaction mixture was prepared per sample: 3.89µl of 2x KASP reaction mix, 0.11µl KASP primers, 1.0µl water and 3.0µl DNA. Plates were then heat sealed and placed into a thermocycler with the following thermal cycling program: 94°C -- 15 minutes, 10 cycles: 94°C -- 20 seconds, 61-53°C (dropping 0.8°C per cycle) -- 60 seconds. 26 cycles: 94°C --20 seconds, 53°C -- 60 seconds, then 10°C until collection.

Following thermal cycling we used a fluorescence reader (Bio-Rad CFX96 Real-Time System) and Bio-Rad CFX Manager software (version 3.1) to automatically determine each samples genotype from a 2d scatter plot of fluorescence in each channel. From this scatter plot outlying samples of unclear genotype were manually excluded from subsequent analysis.

KASP Assay ID: 554-0090.1

KASP Flanking Sequence (alternative allele shown in square brackets, with a forward slash indicating a deletion in the alternative allele): 5’ ACCGCTGGTATGCGATCTGCCACCCGCTAAAATTCAAAAGCACTGCTAAA[A/T]GAGCCCGCAAGAGCATCGTGCTGATCTGGCTGGTGTCCTGCATCATGATG 3’

### Pharmacology

0.15M melatonin and 1M pentylenetetrazole (M5250 and P6500; Sigma, Missouri, USA) stock solutions were made in DMSO and sterile water, respectively. Behavioural testing concentrations for each compound were selected based upon (Rihel *et al.*, 2010). For behaviour experiments each animal in a well with 650µl of fish water was dosed with 1.3µl of either vehicle control or compound at 500x concentration, resulting in a 1 in 500 dilution and thus the desired testing concentration.

### Computing

#### Hardware

A desktop computer with 16GB of RAM was used for most data analysis, figure production and writing. For two-time intensive steps -- hierarchical compression of full module sequences (Batch_Compress.m) and normalising the behavioural motif counts (Batch_Grammar_Freq.m) -- data was run in parallel, with a worker for every animal, on the UCL Legion Cluster (Research Computing Services, UCL).

#### Software

All software used for data handling, analysis and the production of figures is available at https://github.com/ghoshm/Structure_Paper.

#### Processing Behavioural Data

See Supplemental Figure 7 for a flow diagram describing behavioural data acquisition and analysis. All custom behavioural analysis software was written and run in MATLAB 2016b-2018a (MathWorks, Massachusetts, USA). The suffixes .m and .mat denote MATLAB code and MATLAB data files, respectively.

Behavioural data was recorded by subtracting subsequent pairs of frames from each other and determining the number of pixels that changed intensity within each well between each pair of frames, termed Δ pixels. To acquire behaviour data, each Zebrabox was setup using ViewPoint’s ZEBRALAB software (version 3.22), which outputs a .xls and a .raw file (ViewPoint specific format) per experiment. Each behaviour .xls file was reorganised into a .txt file using the function perl_batch_192.m (Jason Rihel). For each experiment a .txt metadata file assigning each animal to an experimental group, for example genotype, was manually produced. To replicate the previous analysis methodology, as in Supplemental Figure 1c, behaviour and metadata .txt files were input to the function sleep_analysis2.m (Jason Rihel).

To assess data on a frame by frame basis, each experiment’s .raw file which was output from ViewPoint’s Zebrabox, was exported within the ZEBRALAB software to thousands of .xls files. Each .xls file contained 50,000 rows and 21 columns, with data from any given well listed approximately every 192 rows, as the setup always assumes recordings are from two 96-well plates. This formatting is, however, only approximate as infrequently the well order is erroneously non-sequential; these rows were termed ordering errors. Each .xls file is formatted with 21 columns, of which 3 contain useful data: type – notes when ViewPoint defined data acquisition errors occurred; location -- denotes which well the data came from; and data1 – records the Δ pixel value from that well for that time point. The function Vp_Extract.m was used to reformat the .xls files from each experiment to single frame by fish matrices, from which each animal’s behaviour was quantified. Vp_Extract.m requires three inputs to be selected: a folder containing the .xls files; a .txt behaviour file output from perl_batch_192.m; and a .txt metadata file. To ensure that each animal has the same number of frames, frames with ViewPoint defined errors or ordering errors (which are automatically detected by Vp_Extract.m) are discarded. A maximum Δ pixels value can be set and active bouts containing even a single frame with a higher Δ pixels value than this are set to zero for the entire duration of the bout. Here a maximum Δ pixels threshold of 200 was set. This value was determined from manual inspection of the dataset as well as by comparisons of this data to data recorded from plates with no animals in. Time periods during which water is being replenished are automatically detected and set to a Δ pixels value of zero. These time periods are noted and excluded from later analysis. The function outputs .mat files for subsequent analysis. Either single or multiple .mat files output from Vp_Extract.m were input to Vp_Analyse.m and Bout_Clustering.m.

Vp_Analyse.m was used to compare general activity levels and bout features across time and between groups. The function has two options. The first allows for specific days and nights of interest to be cropped from the data. The second determines how experimental repeats are handled, treating the data as either a single merged dataset or as separate datasets. In the latter case, each experimental repeat is plotted with the same colour scheme as the first experiment, with progressive shading for each repeat. Additionally, the N-way ANOVA comparisons include a repeat factor, which can be used to determine if results are consistent across experimental repeats. Vp_Analyse.m outputs two statistics results structures: twa -- N-way ANOVA comparison results, and kw -- Two-sample Kolmogorov-Smirnov test results. Vp_Analyse.m outputs figures showing each group’s activity (e.g. Figure 1d-e) and bout features (e.g. Supplemental Figure 2) over time.

The script Bout_Clustering.m was used to cluster all active and inactive bouts into behavioural modules, as well as to compare the resultant modules. To cluster the data an evidence accumulation approach is used (Fred and Jain, 2002, 2005) implemented by the custom MATLAB function gmm_sample_ea.m. Bout_Clustering.m produces figures (e.g. Supplementary Figure 3) and statistically compares the modules. The MATLAB workspace output from Bout_Clustering.m can be input to either Bout_Transitions.m or Bout_Transitions_Hours.m.

The function gmm_sample_ea.m clusters data using an evidence accumulation approach (Fred and Jain, 2002, 2005) through which the results of multiple Gaussian Mixture Models are combined to generate an aggregate solution. This process is executed through the following six steps. Firstly, a sample of ‘probe points’ are randomly sampled from the data. The number of probe points to sample is user defined. Secondly, values of K and sample sizes are uniformly sampled from user set ranges. The values of K are used to set the number of mixture components for each mixture model. The sample sizes determine the number of points, randomly sampled from the data that each mixture model is fit to. Thirdly, a Gaussian Mixture Model is iteratively fit to the sampled data with K components. Each probe point is assigned to the component with the highest corresponding posterior probability and evidence is accumulated on the probe points; evidence is defined as pairwise co-occurrences in the same component. Fourthly, the evidence accumulation matrix is hierarchically clustered, and the final number of clusters is determined by using the maximum differentiated linkage distance to cut the resultant dendrogram. The linkage metric used is a user-defined option. Fifthly, the clusters are normalised for size by randomly sampling the number of points in the smallest cluster, from each cluster. Finally, all data points are assigned to these final size normalized clusters using the mode cluster assignment of the k-nearest neighbours, with k being user defined.

The script Bout_Transitions.m takes the MATLAB workspace output from Bout_Clustering.m as an input and compresses each animal’s full module sequence to generate a library of behavioural motifs. The number of occurrences of each motif are counted and normalised by comparison to paired shuffled data. Finally, a supervised learning algorithm is applied to identify context specific behavioural motifs. For two-time intensive steps -- hierarchical compression of full module sequences (Batch_Compress.m) and normalising the behavioural motif counts (Batch_Grammar_Freq.m) -- data was manually copied (via MobaXterm, Personal Edition v10.5) to UCL Legion Cluster (Research Computing Services, UCL) and processed in parallel with a worker for every fish. MATLAB code for hierarchical compression is described in Gomez-Marin *et al*., (2016). MATLAB code for submitting these jobs to Legion, analysing data and retrieving results is available at https://github.com/ghoshm/Legion_Code. Ultimately, Bout_Transitions.m outputs a library of behavioural motifs and motif related figures (e.g. Figure 3).

The script Bout_Transitions_Hours.m compresses blocks of 500 modules for statistical comparisons, uses the motif library from Bout_Transitions.m to count the occurrence of each motif every hour, normalises these counts to paired shuffled data and finally uses supervised learning to identify hour specific behavioural motifs. As with Bout_Transitions.m behavioural motifs are normalised, via Batch_Grammar_Freq.m, using UCL Legion Cluster. Bout_Transitions_Hours.m outputs figures (e.g. Figure 4d) and statistics.

### Behavioural Data Analysis

#### Δ Pixels

At the acquisition stage, Δ pixels data was filtered by the software (ViewPoint) such that each frame for a given well was scored as either zero or higher. In the absence of movement within a well, and hence no pixels changing intensity, Δ pixels values of zero were recorded. These periods were termed inactive bouts and were defined as any single or consecutive frames with Δ pixels values equal to zero. The length of each inactive bout was used as a descriptive feature. When there was movement within a well, Δ pixels values greater than zero were recorded. These periods were termed active bouts and were defined as any single or consecutive frames with Δ pixels values greater than zero. Six features were used to describe each active bout: length, mean, standard deviation, total, minimum and maximum. These features, as well as the number of active bouts, percentage of time spent active and total Δ pixels activity, were compared between conditions, e.g. day and night and dose of drug, in two ways using the function Vp_Analyse.m.

To compare the distribution of values for each feature between conditions, a probability density function (pdf) was fit to each animal’s data and the mean shape of each condition’s pdf was compared using a Two-sample Kolmogorov-Smirnov test (e.g. Supplementary Figure 2a). To compare each feature’s average values between conditions, mean values were taken from each animal, and N-way analysis of variance was computed. The following factors, when relevant, were included and full interaction terms were calculated: condition -- e.g. mutant and wild-type; time -- e.g. day and night; development -- defined as a consecutive day and night; and experimental repeat -- i.e. which experimental repeat a datapoint came from. For experiments with multiple repeats, the lack of an interaction effect between the comparison of interest and experimental repeat factor was considered as evidence of a consistent result.

Note that our analysis relies on periods of inactivity registering Δ pixels values of zero. Consequently, to be compatible with our analysis code, other data may require post-recording filtering.

#### Clustering

To cluster the bouts, the script Bout_Clustering.m was used. First, matrices of bouts by features were constructed (Active matrix -- 30,900,018 x 6; Inactive matrix -- 30,900,418 x 1). To prepare the active data for clustering each animal’s data was individually normalised by calculating z-scores using equation 1, which illustrates how every bout (i) from each animal (f) was normalised by first subtracting the mean of this animal’s bout features (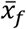) from the bout and then dividing by the standard deviation of each bout feature for this animal σ_f_.

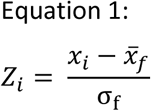

Active bout features across all animals were then centred by subtracting each feature’s mean value from every bout, and principal component analysis (PCA) was used to reduce the data to 3 dimensions, the knee point of the scree plot, which together explain 97.5% of the variance (Supplementary Figure 3a).

Next, the active and inactive bouts were separately clustered using an evidence accumulation-based approach (Fred and Jain, 2002, 2005) implemented by the function gmm_sample_ea.m. Firstly, 40,000 probe points were randomly sampled from the data. Next, for 200 iterations, another group of points were randomly sampled and fit with a Gaussian mixture model with a random number of clusters. For each iteration, these two parameters varied uniformly in the following ranges: the number of points sampled -- 40,000 to 100,000; the number of clusters fit -- 2 to 20. Each mixture model was fit using MATLAB’s fitgmdist function (MATLAB, Statistics and Machine Learning Toolbox) with full, regularized, independent covariance matrices and initialised using the k-means++ algorithm (Arthur and Vassilvitskii, 2007). Each mixture model was fit 5 times and the one with the largest log-likelihood was retained. Once each model had been fit, each probe point was assigned to the component with the largest posterior probability, and evidence in the form of pairwise occurrence in the same cluster was accumulated on the probe points. Once the 200 mixture models had been fit, average link clustering was applied to the evidence accumulation matrix and the final number of clusters determined based on maximum cluster lifetime. Next, the resultant clusters were normalised for size by randomly selecting the number of points in the smallest cluster from each cluster (5,983 active, 614 inactive bouts). Finally, all points were assigned to the size normalised clusters using the mode cluster assignment of the 50 nearest neighbours for every point.

#### Hierarchical Compression

Clustering reduced each animal’s behaviour to a non-repetitive sequence of active and inactive bouts, termed modules. On average this reduced each wild-type sequence length by 96%, from 6,308,514 frames to 236,636 modules, easing the computational demands of compressing these sequences.

To compress modular sequences, an offline compressive heuristic (Nevill-Manning and Witten, 2000) was used (equation 2). At each iteration (i) of the algorithm, the most compressive motif was defined as the motif which made the most savings, a balance between the length of the motif (W) and the number of times it occurred in the sequence (N), which also considered the combined cost of adding a new motif to the dictionary (W + 1) and of introducing a new symbol into the sequence (+N) at every occurrence of this motif in the sequence.

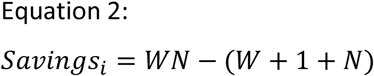

The overall compressibility of a given input sequence was calculated by summing these savings across all iterations and dividing this total by the length of the original input sequence (in modules). This process resulted in a compressibility metric that ranged from 0-1 (low-high compressibility). To reduce computational time, motifs of a maximum of 10 modules long were sought, although the hierarchical nature of the algorithm enabled the identification of longer motifs through nesting. To generate the common motif library, the motifs obtained from compression of every animal’s full module sequence (Batch_Compress.m) were merged, and then all unique motifs were kept (Bout_Transitions.m). To generate sets of paired control sequences for every animal, each animal’s module sequence was divided into sequential day and night or hourly segments and the modules within each of these windows was shuffled 10 times, maintaining the active/inactive transition structure (Bout_Transitions.m). As compressibility varies non-linearly with uncompressed sequence length (Supplementary Figure 5b), to enable comparisons between samples with different numbers of modules, non-overlapping blocks 500 modules long were compressed (Bout_Transitions_Hours.m).

#### Supervised Motif Selection

To identify both which and how many motifs were required to distinguish between behavioural contexts (e.g. day and night), the following approach was executed by the function Batch_Grammar_Freq.m. Firstly, the number of occurrences of every motif from the common motif library was counted in every real and shuffled modular sequence. Next, to calculate enrichment/constraint scores for every motif, the deviation of the real from shuffled counts, as well as the deviation of each shuffle from the other shuffles, was calculated (equation 3). For a given animal and time window, i.e. day or night, the mean number of times motif (i) was counted in the shuffled data (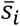), was subtracted from the real number of counts (*x*_*i*_) and divided by the standard deviation of the shuffled counts (*σ*_*si*_).

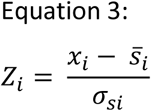

When comparing the shuffled data to itself, each shuffle (now *x*_*i*_) was excluded from 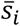 and *σ*_*si*_. Infinite values occurred when there was no standard deviation in the *σ*_*si*_ counts and thus *σ*_*si*_ equalled zero. For subsequent working, infinite values were replaced with a constant value of ± 3.32. This value was chosen as equation 3 will always output this value when there is no standard deviation in the shuffled counts and *x*_*i*_ is included in the calculation of *σ*_*si*_. Note that in the real data, infinite values constituted only 2.2% of all enrichment/ constraint scores.

For any given comparison, motif library enrichment/constraint scores for the relevant animals were formatted into a matrix of samples by motifs (e.g. Figure 4b). Scores for each motif (column) were normalised by subtracting each column’s mean score and dividing by each column’s standard deviation. A supervised feature selection algorithm (Peng *et al.*, 2005) was applied to these matrices to select the top 250 maximally relevant and minimally redundant (mRMR) motifs. To determine how many of these motifs were necessary for accurate classification, linear discriminant analysis classifiers were trained on this data using 10-fold cross validation as sequential mRMR motifs were added, and classification error mean and standard deviation were calculated. The MATLAB function fitcdiscr (Statistics and Machine Learning Toolbox) was used to implement these steps. Finally, to determine how many motifs were necessary for a given comparison, classification error curves were smoothed with a running average 3 motifs wide and the number of motifs at which the minimum classification error occurred was identified (Supplementary Figure 6a). To evaluate classifier performance, the results of each classifier were compared to a majority class classifier whose performance depended upon the ratio of samples of each class. For example, in a dataset with two labels at a ratio of 0.1 : 0.9, the majority class classifier would consistently assign the latter label and achieve a classification error of 10% (± standard error of proportion). Additionally, we compared each classifiers performance to a set of 10 classifiers built using the same number of motifs, though randomly selected. For example, for a classifier which achieved its minimal classification error using 15 motifs, we randomly selected 15 motifs and trained classifiers as above. We repeated this process 10 times per classifier and report the error and standard deviation across these 10 repeats.

**Table 1.**
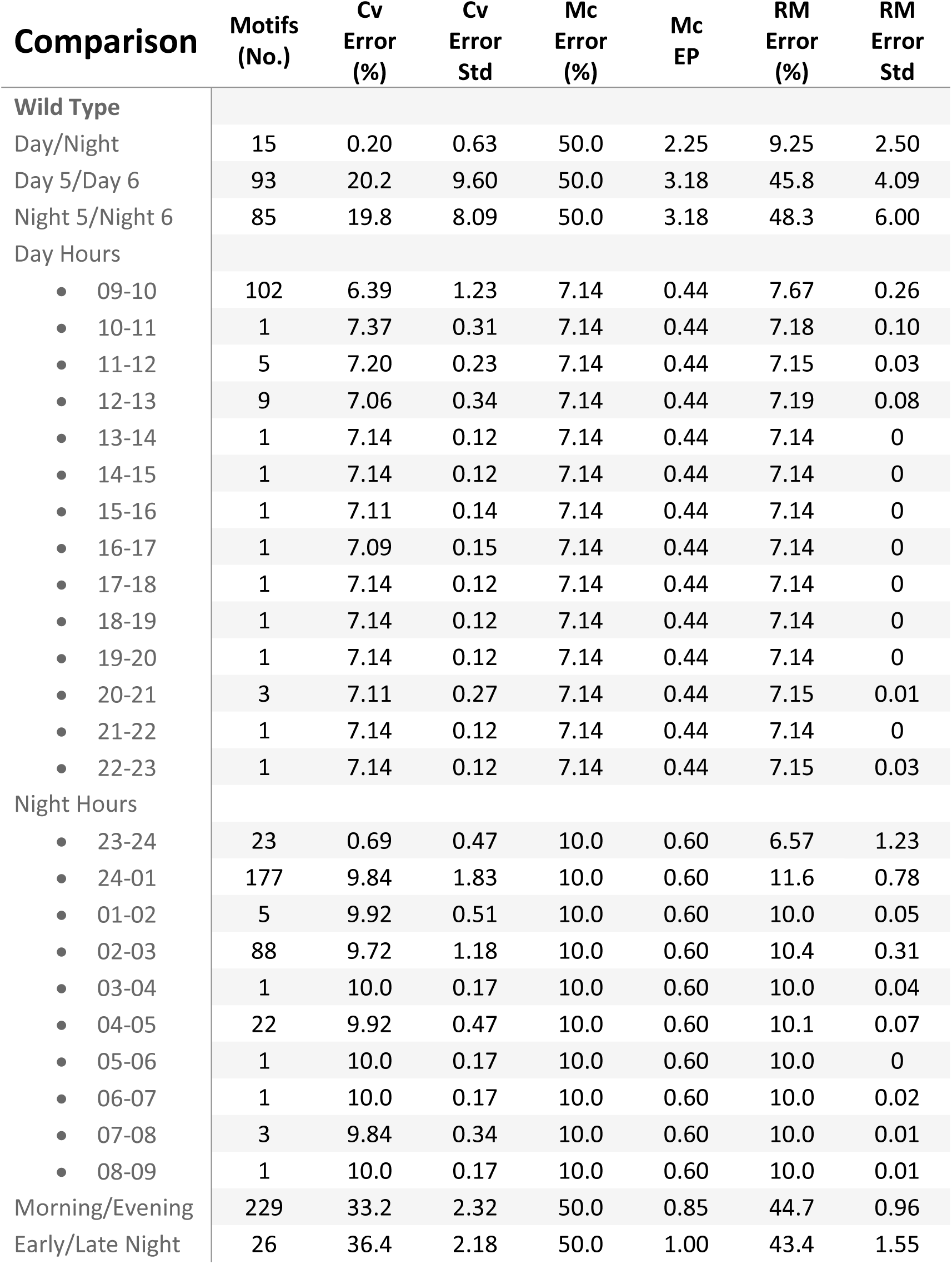
Wild-type Motif Classifier Performance. A table showing the performance of each wild-type motif classifier. Each classifier sought to separate the data shown in the comparison column, e.g. Day/Night. For the hourly comparisons, each hour was compared to data from all other hours grouped together. For each comparison 250 motifs were chosen by mRMR, then a smaller number were retained (see Motifs column) based on classification error curves (see Supplementary Figure 6a). No. – number. Cv – 10-fold cross validated. Std – standard deviation across the 10 folds. Mc – majority class classifier. EP – standard error of proportion. RM – classifiers built from random motif subsets.

| Comparison | Motifs<br>(No.) | Cv<br>Error<br>(%) | Cv<br>Error<br>Std | Mc<br>Error<br>(%) | Mc<br>EP | RM<br>Error<br>(%) | RM<br>Error<br>Std |
| --- | --- | --- | --- | --- | --- | --- | --- |
| **Wild Type** |  |  |  |  |  |  |  |
| Day/Night | 15 | 0.20 | 0.63 | 50.0 | 2.25 | 9.25 | 2.50 |
| Day 5/Day 6 | 93 | 20.2 | 9.60 | 50.0 | 3.18 | 45.8 | 4.09 |
| Night 5/Night 6 | 85 | 19.8 | 8.09 | 50.0 | 3.18 | 48.3 | 6.00 |
| Day Hours |  |  |  |  |  |  |  |
| • 09-10 | 102 | 6.39 | 1.23 | 7.14 | 0.44 | 7.67 | 0.26 |
| • 10-11 | 1 | 7.37 | 0.31 | 7.14 | 0.44 | 7.18 | 0.10 |
| • 11-12 | 5 | 7.20 | 0.23 | 7.14 | 0.44 | 7.15 | 0.03 |
| • 12-13 | 9 | 7.06 | 0.34 | 7.14 | 0.44 | 7.19 | 0.08 |
| • 13-14 | 1 | 7.14 | 0.12 | 7.14 | 0.44 | 7.14 | 0 |
| • 14-15 | 1 | 7.14 | 0.12 | 7.14 | 0.44 | 7.14 | 0 |
| • 15-16 | 1 | 7.11 | 0.14 | 7.14 | 0.44 | 7.14 | 0 |
| • 16-17 | 1 | 7.09 | 0.15 | 7.14 | 0.44 | 7.14 | 0 |
| • 17-18 | 1 | 7.14 | 0.12 | 7.14 | 0.44 | 7.14 | 0 |
| • 18-19 | 1 | 7.14 | 0.12 | 7.14 | 0.44 | 7.14 | 0 |
| • 19-20 | 1 | 7.14 | 0.12 | 7.14 | 0.44 | 7.14 | 0 |
| • 20-21 | 3 | 7.11 | 0.27 | 7.14 | 0.44 | 7.15 | 0.01 |
| • 21-22 | 1 | 7.14 | 0.12 | 7.14 | 0.44 | 7.14 | 0 |
| • 22-23 | 1 | 7.14 | 0.12 | 7.14 | 0.44 | 7.15 | 0.03 |
| Night Hours |  |  |  |  |  |  |  |
| • 23-24 | 23 | 0.69 | 0.47 | 10.0 | 0.60 | 6.57 | 1.23 |
| • 24-01 | 177 | 9.84 | 1.83 | 10.0 | 0.60 | 11.6 | 0.78 |
| • 01-02 | 5 | 9.92 | 0.51 | 10.0 | 0.60 | 10.0 | 0.05 |
| • 02-03 | 88 | 9.72 | 1.18 | 10.0 | 0.60 | 10.4 | 0.31 |
| • 03-04 | 1 | 10.0 | 0.17 | 10.0 | 0.60 | 10.0 | 0.04 |
| • 04-05 | 22 | 9.92 | 0.47 | 10.0 | 0.60 | 10.1 | 0.07 |
| • 05-06 | 1 | 10.0 | 0.17 | 10.0 | 0.60 | 10.0 | 0 |
| • 06-07 | 1 | 10.0 | 0.17 | 10.0 | 0.60 | 10.0 | 0.02 |
| • 07-08 | 3 | 9.84 | 0.34 | 10.0 | 0.60 | 10.0 | 0.01 |
| • 08-09 | 1 | 10.0 | 0.17 | 10.0 | 0.60 | 10.0 | 0.01 |
| Morning/Evening | 229 | 33.2 | 2.32 | 50.0 | 0.85 | 44.7 | 0.96 |
| Early/Late Night | 26 | 36.4 | 2.18 | 50.0 | 1.00 | 43.4 | 1.55 |

**Table 2.** *hcrtr* and Pharmacological Classifier Performance. A table showing the performance of each classifier. Each classifier sought to separate the data shown in the comparison column, e.g. *hcrtr*^+/+^ (WT) and *hcrtr*^-/+^ (Het). For the pharmacological comparisons each condition was compared to the rest of the conditions grouped together, aside from the control data which was excluded. For each comparison 250 motifs were chosen by mRMR, then a smaller number were retained (see Motifs column) based on classification error curves (see Supplementary figure 6a). No. – number. Cv – 10-fold cross validated. Std – standard deviation across the 10 folds. Mc – majority class classifier. EP – standard error of proportion. RM – classifiers built from random motif subsets. WT - *hcrtr*^+/+^, Het – *hcrtr*^-/+^, Hom – *hcrtr*^-/-^.

| Comparison | Motifs<br>(No.) | Cv<br>Error<br>(%) | Cv<br>Error<br>Std | Mc<br>Error<br>(%) | Mc<br>EP | RM<br>Error<br>(%) | RM<br>Error<br>Std |
| --- | --- | --- | --- | --- | --- | --- | --- |
| *hcrtr* |  |  |  |  |  |  |  |
| Day and Night |  |  |  |  |  |  |  |
| • WT/Het | 173 | 25.5 | 6.77 | 27.7 | 1.88 | 39.0 | 1.93 |
| • WT/Hom | 83 | 24.7 | 6.07 | 50.0 | 2.83 | 47.8 | 3.66 |
| • Het/Hom | 235 | 24.7 | 3.76 | 27.7 | 1.88 | 38.6 | 1.83 |
| Day |  |  |  |  |  |  |  |
| • WT/Het | 80 | 19.5 | 9.60 | 27.7 | 2.66 | 37.9 | 1.08 |
| • WT/Hom | 195 | 16.7 | 7.50 | 50.0 | 4.00 | 48.7 | 4.22 |
| • Het/Hom | 55 | 22.7 | 7.02 | 27.7 | 2.66 | 33.8 | 2.01 |
| Night |  |  |  |  |  |  |  |
| • WT/Het | 79 | 16.3 | 6.38 | 27.7 | 2.66 | 37.1 | 7.17 |
| • WT/Hom | 53 | 12.8 | 9.58 | 50.0 | 4.00 | 52.3 | 6.16 |
| • Het/Hom | 76 | 16.0 | 7.27 | 27.7 | 2.66 | 36.0 | 5.05 |
| Melatonin (Day) |  |  |  |  |  |  |  |
| • Control | 40 | 0 | 0 | 25.0 | 4.42 | 16.4 | 3.67 |
| • 0.01μM | 89 | 1.39 | 4.52 | 16.7 | 4.39 | 30.0 | 15.2 |
| • 0.1μM | 192 | 1.39 | 4.52 | 16.7 | 4.39 | 20.5 | 18.0 |
| • 1μM | 132 | 2.78 | 6.02 | 16.7 | 4.39 | 29.5 | 8.67 |
| • 3μM | 97 | 0 | 0 | 16.7 | 4.39 | 48.6 | 11.2 |
| • 10μM | 250 | 2.78 | 6.02 | 16.7 | 4.39 | 20.0 | 9.40 |
| • 30μM | 133 | 0 | 0 | 16.7 | 4.39 | 32.2 | 7.89 |
| PTZ (Day) |  |  |  |  |  |  |  |
| • Control | 26 | 0 | 0 | 46.2 | 6.91 | 15.8 | 6.43 |
| • 2.5mM | 55 | 0 | 0 | 35.7 | 9.06 | 42.1 | 11.4 |
| • 5mM | 162 | 0 | 0 | 32.1 | 8.83 | 34.3 | 18.0 |
| • 7.5mM | 104 | 0 | 0 | 32.1 | 8.83 | 49.9 | 14.7 |

**Table 3.** Module Classifier Performance. A table showing the performance of each module classifier. Each classifier sought to separate the data shown in the comparison column, e.g. Wild Type, Day/Night. For each comparison all 10 modules were sequentially chosen by the mRMR algorithm, then a smaller subset was retained (see Module column) based on classification error curves. No. – number. Cv – 10-fold cross validated. Std – standard deviation across the 10 folds. Mc – majority class classifier. EP – standard error of proportion.

| Comparison | Modules<br>(No.) | Cv<br>Error<br>(%) | Cv<br>Error<br>Std | Mc<br>Error<br>(%) | Mc<br>EP |
| --- | --- | --- | --- | --- | --- |
| **Wild Type** |  |  |  |  |  |
| • Day/Night | 10 | 1.61 | 1.29 | 50.0 | 2.25 |
| • Day 5/Day 6 | 8 | 21.0 | 6.53 | 50.0 | 3.18 |
| • Night 5/Night 6 | 1 | 35.5 | 9.71 | 50.0 | 3.18 |
| **hcrtr** |  |  |  |  |  |
| **Day and Night** |  |  |  |  |  |
| • WT/Het | 1 | 27.7 | 0.77 | 27.7 | 1.88 |
| • WT/Hom | 10 | 45.8 | 10.9 | 50.0 | 2.83 |
| • Het/Hom | 8 | 27.5 | 1.12 | 27.7 | 1.88 |
| **Day** |  |  |  |  |  |
| • WT/Het | 1 | 27.7 | 1.46 | 27.7 | 2.66 |
| • WT/Hom | 1 | 40.4 | 12.5 | 50.0 | 4.00 |
| • Het/Hom | 3 | 27.3 | 2.35 | 27.7 | 2.66 |
| **Night** |  |  |  |  |  |
| • WT/Het | 1 | 27.7 | 1.46 | 27.7 | 2.66 |
| • WT/Hom | 1 | 47.4 | 10.9 | 50.0 | 4.00 |
| • Het/Hom | 10 | 27.0 | 1.72 | 27.7 | 2.66 |
| **Melatonin (Day)** |  |  |  |  |  |
| • Control | 3 | 8.33 | 8.69 | 25.0 | 4.42 |
| • 0.01μM | 10 | 2.78 | 6.02 | 16.7 | 4.39 |
| • 0.1μM | 2 | 16.7 | 4.52 | 16.7 | 4.39 |
| • 1μM | 1 | 18.1 | 7.74 | 16.7 | 4.39 |
| • 3μM | 1 | 16.8 | 8.67 | 16.7 | 4.39 |
| • 10μM | 1 | 16.8 | 4.52 | 16.7 | 4.39 |
| • 30μM | 1 | 16.8 | 4.52 | 16.7 | 4.39 |
| **PTZ (Day)** |  |  |  |  |  |
| • Control | 1 | 1.92 | 5.27 | 46.2 | 6.91 |
| • 2.5mM | 1 | 17.9 | 17.6 | 35.7 | 9.06 |
| • 5mM | 1 | 28.6 | 22.3 | 32.1 | 8.83 |
| • 7.5mM | 10 | 20.0 | 26.1 | 32.1 | 8.83 |

## Supplementary Information

### Data Sharing

Processed behavioural data is available at: https://zenodo.org/record/3344770#.XYYwYyhKiUk. Raw data is available upon request.

### Supplementary video 1. High-throughput Behavioural Tracking

A video of 96, 6dpf zebrafish larvae swimming in our rig. The last 1 second of each larva’s Δ pixels data is plotted over each well. This video was filmed at 25Hz and is played back in real time.

### Supplementary video 2. Behavioural Modules

A video of 96, 6dpf zebrafish larvae swimming in our rig. The last 1 second of each larva’s Δ pixels data is plotted over each well, with each active and inactive bout coloured according to its module assignment. This video was filmed at 25Hz and is played back in real time.

## Acknowledgements

We thank members of the UCL fish floor for insightful discussions, Ida Barlow and François Kroll for comments on the manuscript, and the UCL Fish Facility and staff for support.

## Author Contributions

M.G. and J.R. conceived the experiments. M.G. performed the experiments. M.G. designed, wrote code to, and executed data handling and analysis. M.G. produced and formatted figures. M.G. and J.R. wrote the paper.

---

**Supplementary Figure 1.**
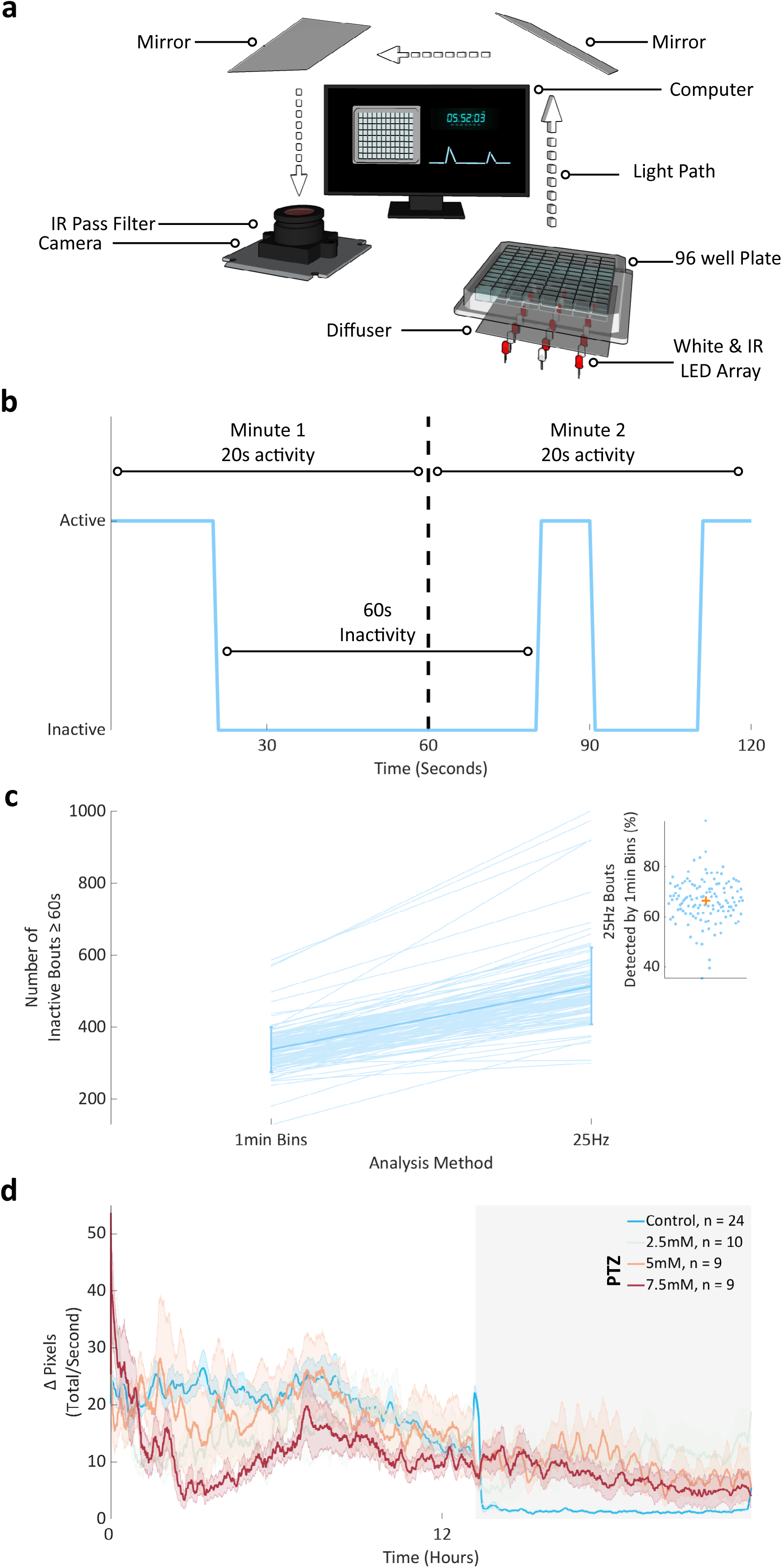
Behavioural Set-up and Analysis. **a.** Schematic of our behavioural set-up. Note that aside from the computer, the set-up is fully enclosed. Not shown to scale. IR - infra-red, LED - light emitting diode. **b.** A fictive illustration of zebrafish behaviour (blue line). Two minutes of data are shown divided by a black dashed vertical line. A 1min binning approach would score both minutes as 20 seconds of activity and miss the 60 second period of inactivity in between. This latter loss leads to a discrepancy in the number of periods ≥ 60s between the 1-minute bin and 25Hz methods (see c). **c.** The number of inactive periods ≥ 60s for each of 124 wild type animals is shown, as determined by both a 1-minute bin and 25Hz approach. Data is from each animal’s entire recording period (4-7dpf). Data for each animal is shown as a pale blue line overlaid with a bold line showing the population mean and standard deviation. Insert: the percentage of the 25Hz counts detected by the 1minute bin method per animal. Each animal’s data is shown by a circle. An orange cross marks the population mean. **d.** Average activity across one day (white background) and night (dark background) for larvae exposed to either H_2_O (control) or a range of PTZ doses immediately prior to tracking at 6dpf. Data for each larva was summed into seconds and then smoothed with a 15-minute running average. Shown is a mean summed and smoothed trace (bold line) and standard error of the mean (shaded surround). n denotes the number of animals per condition.

**Supplementary Figure 2.**
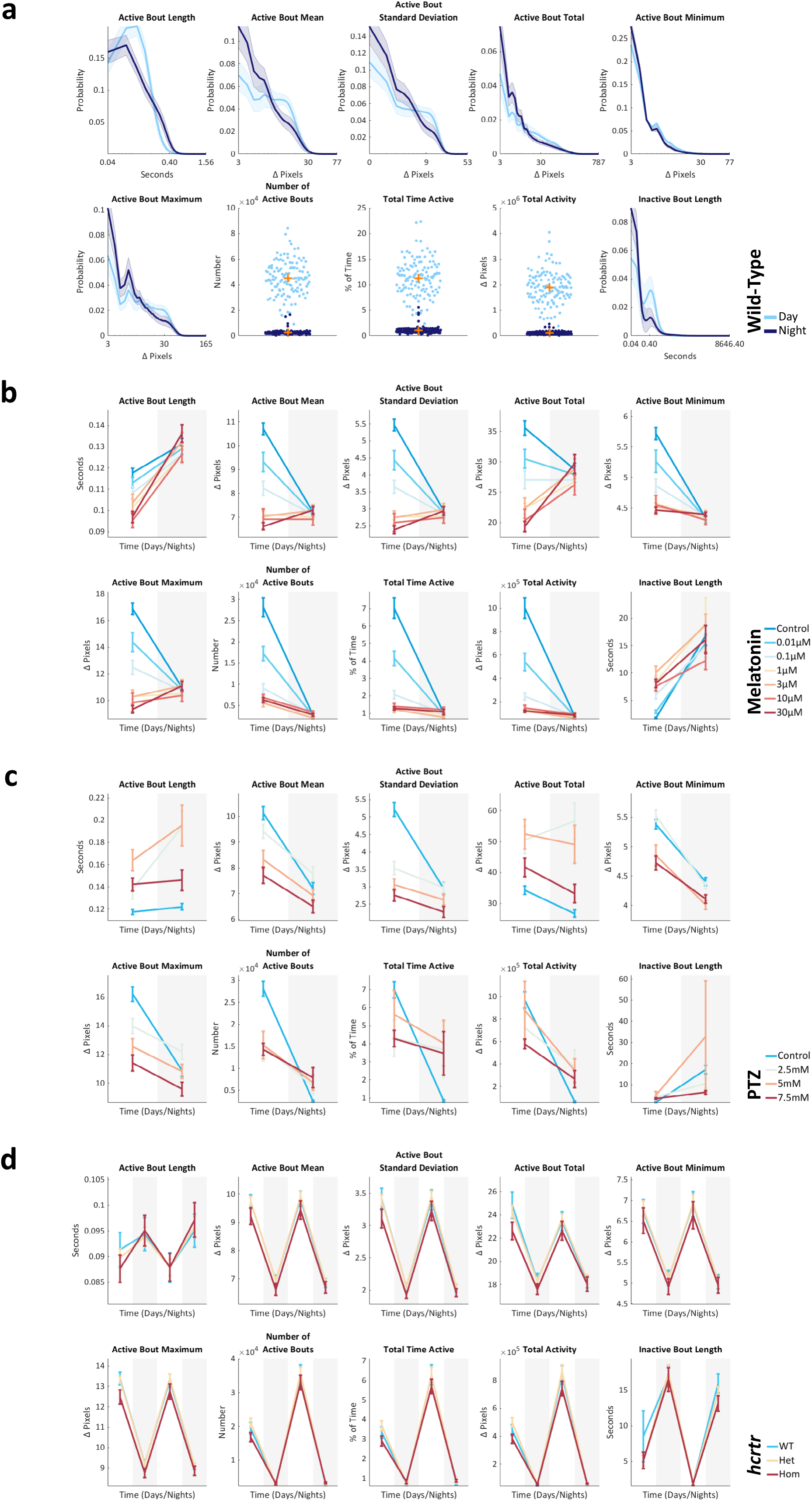
Bout Features. **a.** Bout feature distributions during the day (light blue) and the night (dark blue). For the probability curves, each animal’s data was fit with a probability density function (pdf). Shown is a mean pdf (bold line) and standard deviation (shaded surround) with a log scale on the x-axis. For the scatter plots, each larva’s mean value across the days or nights (5-6dpf) is shown as a light blue (day) or dark blue circle (night). An orange cross marks each population’s mean. Of the pdfs, only the mean day and night active bout total and inactive bout length pdfs were consistently significantly different across three independent experiments (p < 0.01; Two-sample Kolmogorov-Smirnov test). n = 124 wild-type larvae. **b.** Melatonin bout feature means. A mean was taken per animal per feature, and day or night (6dpf). Shown is a population mean and standard error of the mean during the day (white background) and the night (grey background). Control - DMSO. n = 24 controls then n = 12 per dose. **c.** PTZ bout feature means, as in b. Control - H_2_O. n = 24 controls then n = 10 (2.5mM), n = 9 (5mM) and n = 9 (7.5mM). **d.** *hcrtr* bout feature means as in b, for days (white background) and nights (grey background) 5 to 6 post fertilisation. *hcrtr*^-/-^ mutants had significantly lower mean values compared to both *hcrtr*^+/+^ and *hcrtr*^-/+^ for the following active bout features: length, standard deviation and total (p < 0.05 for all comparisons, Dunn-Sidak corrected four-way ANOVA, adjusted for the following factors: day/night, development and experimental repeat). No features differed significantly between *hcrtr*^-/+^ and *hcrtr*^+/+^. n = 39, 102 and 39, for WT - *hcrtr*^+/+^, Het – *hcrtr*^-/+^, Hom – *hcrtr*^-/-^ respectively.

**Supplementary Figure 3.**
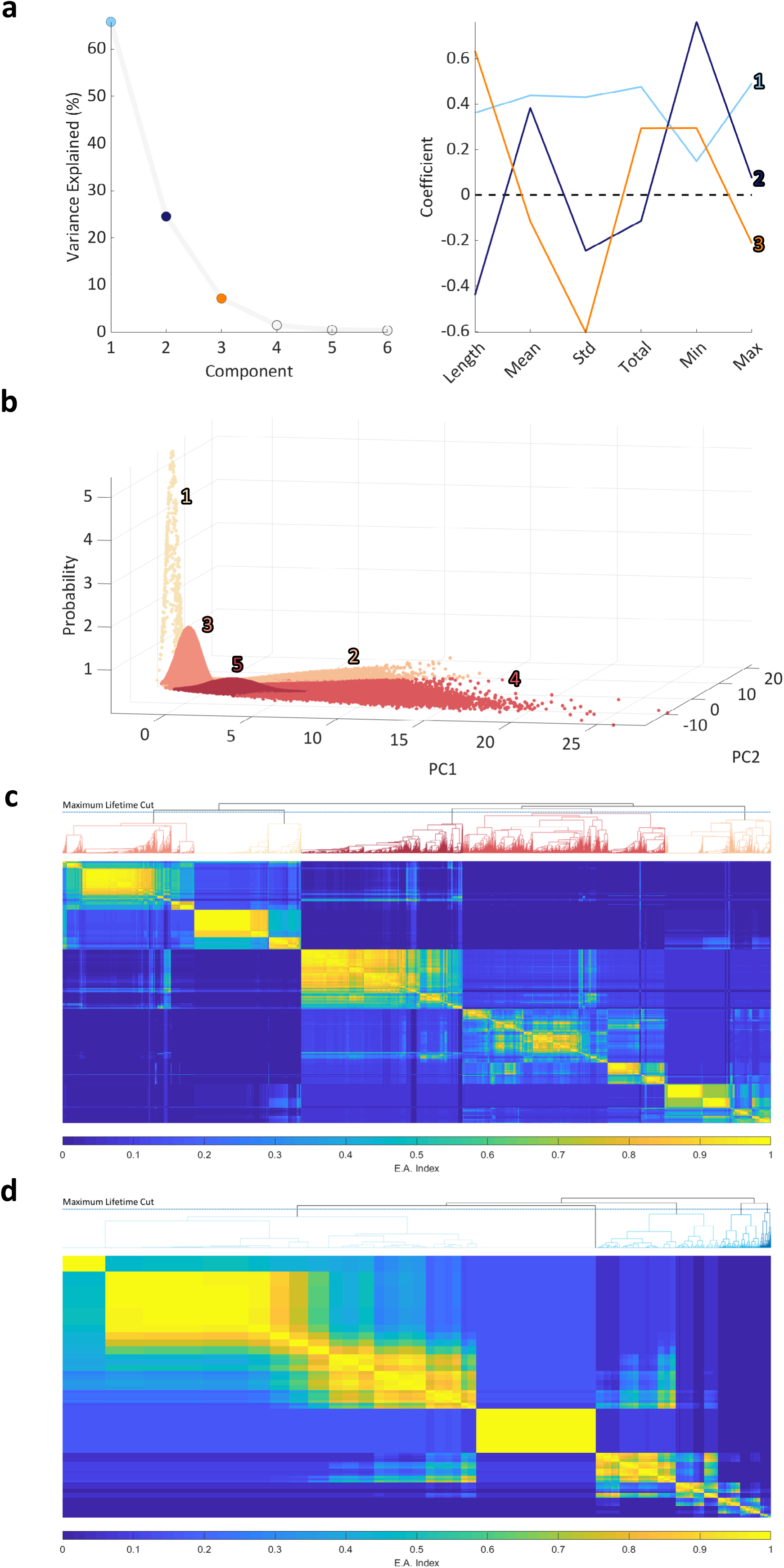
Evidence-accumulation Based Clustering. **a.** Left: scree plot showing the percentage of variance explained by each principal component from the active bout data. The first 3 principal components, the knee point of the curve, were kept for subsequent analysis. The colours of these points refer to the right panel. Right: each of the 3 retained component’s coefficients for the different active bout parameters is shown. **b.** The active bouts within each module were fit by Gaussian distributions. Each active bout is shown in a 3D space of PC1, PC2, and probability. Each bout is numbered and coloured by its module assignment. **c.** Evidence accumulation (E.A.) matrix for the 40,000 active probe points (matrix dimensions are thus 40,000 by 40,000). A higher E.A. index indicates a higher frequency of pairwise occurrences in the same cluster across 200 Gaussian Mixture Models. This matrix was clustered hierarchically, and a maximum lifetime cut was made to determine the final number of modules. The dendrogram above shows all 40,000 leaves and is coloured by mean module length from shortest (lightest) to longest (darkest) as in other figures. **d.** Evidence accumulation matrix for the inactive bouts.

**Supplementary Figure 4.**
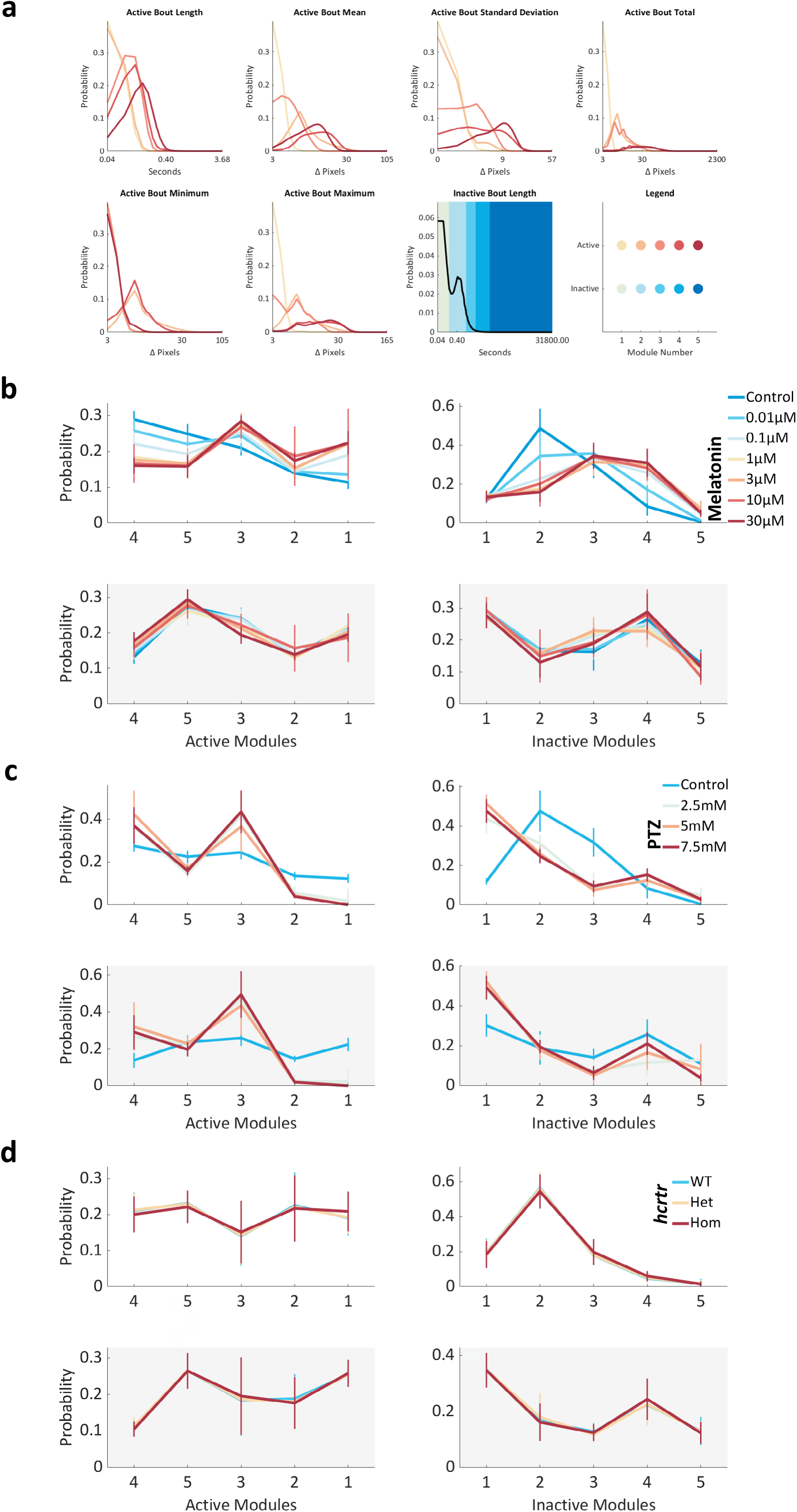
Behavioural Modules. **a.** Probability density functions for each bout feature by module. All features are shown on a log x-axis. The legend panel indicates each module’s colour. **b.** Melatonin module probabilities during 6dpf day (upper panels) and night (lower panels) for both the active (left) and inactive (right) modules. Shown is a mean and standard error of the mean for each group, coloured according to the legend. Active modules are sorted from highest to lowest by average wild type day probability, based upon wild type data in Figure 2d. Inactive modules are sorted by increasing mean length. Control - DMSO. n = 24 controls then n = 12 per dose. **c.** PTZ data as in b, with H_2_O (control). n = 24 controls then n = 10 (2.5mM), n = 9 (5mM) and n = 9 (7.5mM). **d.** *hcrtr* data as in b, with mean values across 5 and 6dpf. No module probabilities differed significantly among genotypes (full four-way ANOVA, with the following factors: genotype, day/night, development, and experimental repeat). n = 39, 102 and 39, for WT - *hcrtr*^+/+^, Het – *hcrtr*^-/+^, Hom – *hcrtr*^-/-^ respectively.

**Supplementary Figure 5.**
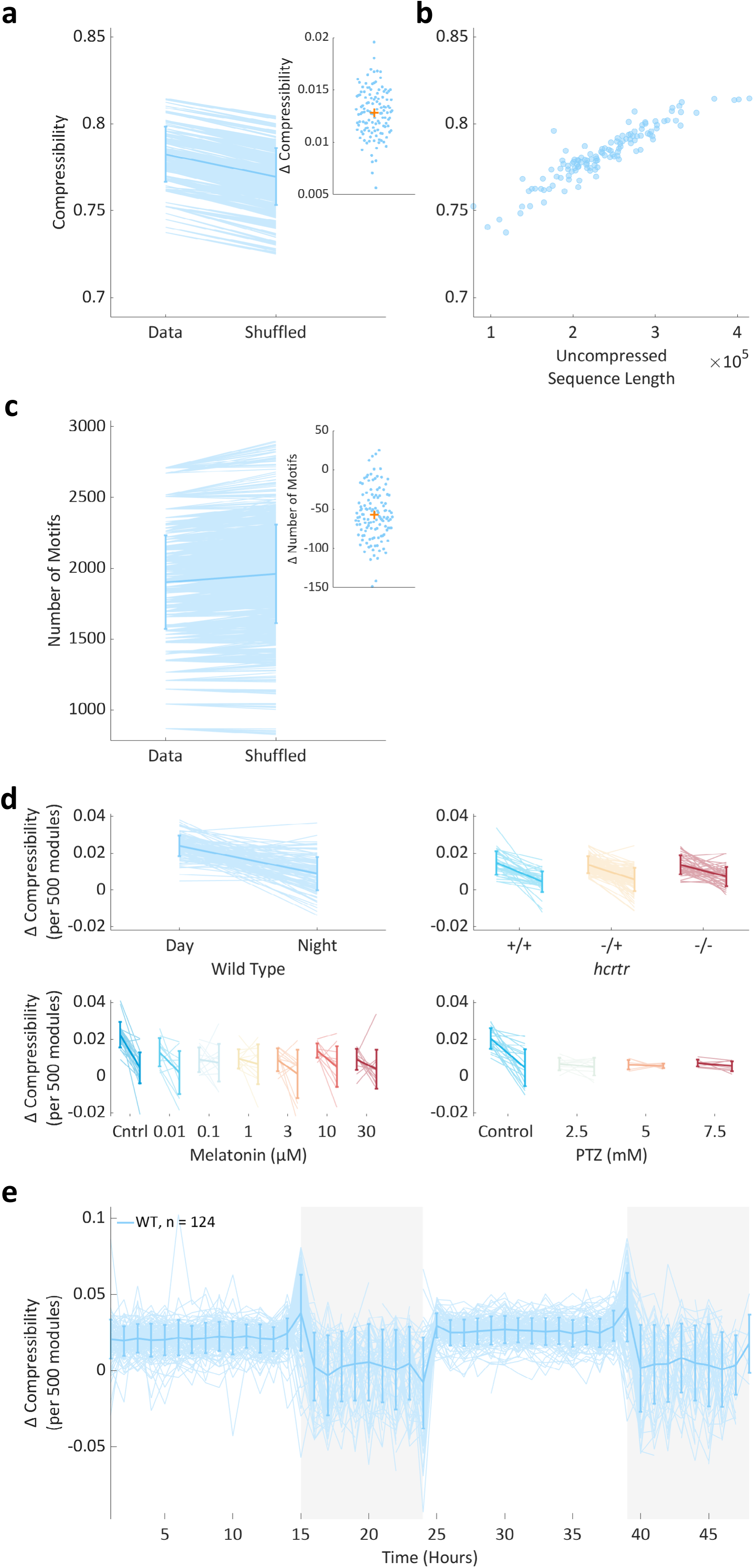
Hierarchical Compression Metrics. **a.** The compressibility (y-axis) of the real wild-type data is higher than the paired shuffled data (p < 10^-15^, two-way ANOVA, real vs shuffled data, no significant interaction with experimental repeat factor). Each animal’s data is shown as a pale blue line. Overlaid is a mean and standard deviation. Insert: the mean difference in compressibility between each larva’s real and shuffled data. Each larva is shown by a circle, and the orange cross marks the mean. **b.** The compressibility (y-axis) of the real wild type data varies non-linearly with uncompressed sequence length. Each larva (of 124) is shown as a dot. **c.** The number of motifs (y-axis) identified from compressing each wild-type animal’s real and paired shuffled data. Each animal’s data is shown as a pale blue line. Overlaid is a mean and standard deviation. Insert: the mean intra-fish difference in the number of identified motifs. Each larva is shown by a circle, and the orange cross marks the mean. **d.** Each panel shows how Δ compressibility varies in different behavioural contexts. Each pale line shows an individual larva’s average Δ compressibility during the day and the night. The darker overlay shows a population day and night mean and standard deviation. **e.** Δ Compressibility of 500 module blocks for each wild-type larva, averaged into 1-hour time points. Each pale blue line shows 1 of 124 larvae. Line breaks occur when a larva had less than 500 modules within a given hour. The darker blue overlay shows the mean and standard deviation of this data every hour. Shown are days (white background) and nights (dark background) 5 and 6 of development.

**Supplementary Figure 6.**
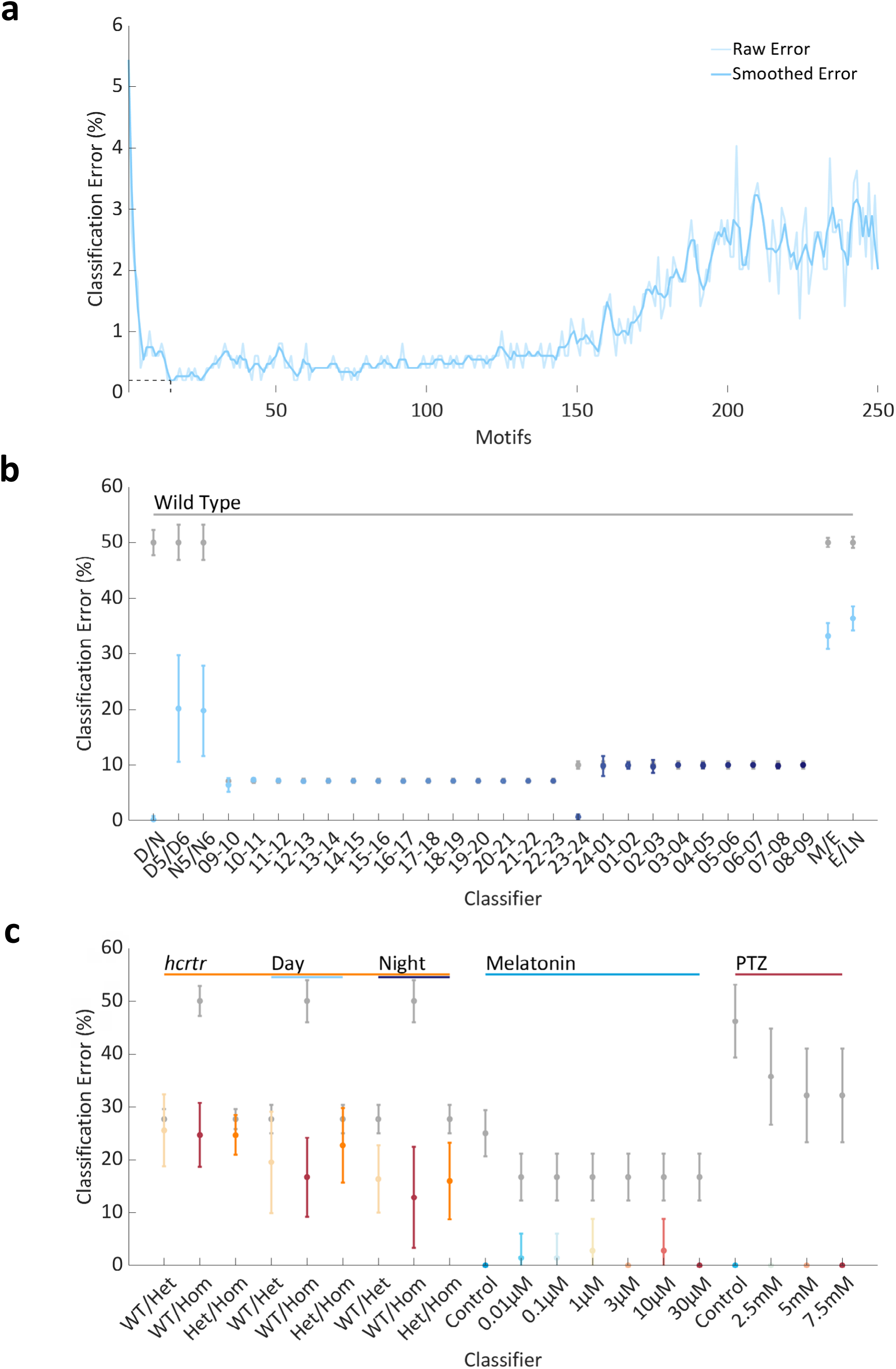
Motif Classifier Performance. **a.** Classification error (%) from linear classifiers separating wild-type day and night behaviour using motif enrichment/constraint scores as sequential mRMR motifs from 1-250 are added (x-axis). The average error is shown in light blue. Overlaid in darker blue is a running average 3 motifs wide. The broken black lines show the minimum of the smoothed data to be at 15 motifs, where the classification error is 0.2%. **b.** Wild-type temporal classifier performance. Real classifiers (colour) are shown as a mean and standard deviation from 10-fold cross validation. Majority class classifiers (grey) are shown as value and standard error of proportion. Each classifiers data is listed on the x-axis. D - day, N - night, M/E - morning/evening, E/LN - early/late night. The number of motifs chosen for each classification and exact values for each classifier are detailed in Supplementary Table 1. **c.** *hcrtr*, Melatonin and PTZ classifier performance. Real classifiers (colour) are shown as a mean and standard deviation from 10-fold cross validation. Majority class classifiers (grey) are shown as value and standard error of proportion. Each classifier’s data is listed on the x-axis. For *hcrtr* comparisons, grouped classifiers as well as separate day (light blue underline) and night (dark blue underline) classifiers are shown. For melatonin and PTZ, only day data was compared. Classifier details can be found in Supplementary Table 2.

**Supplementary Figure 7.**
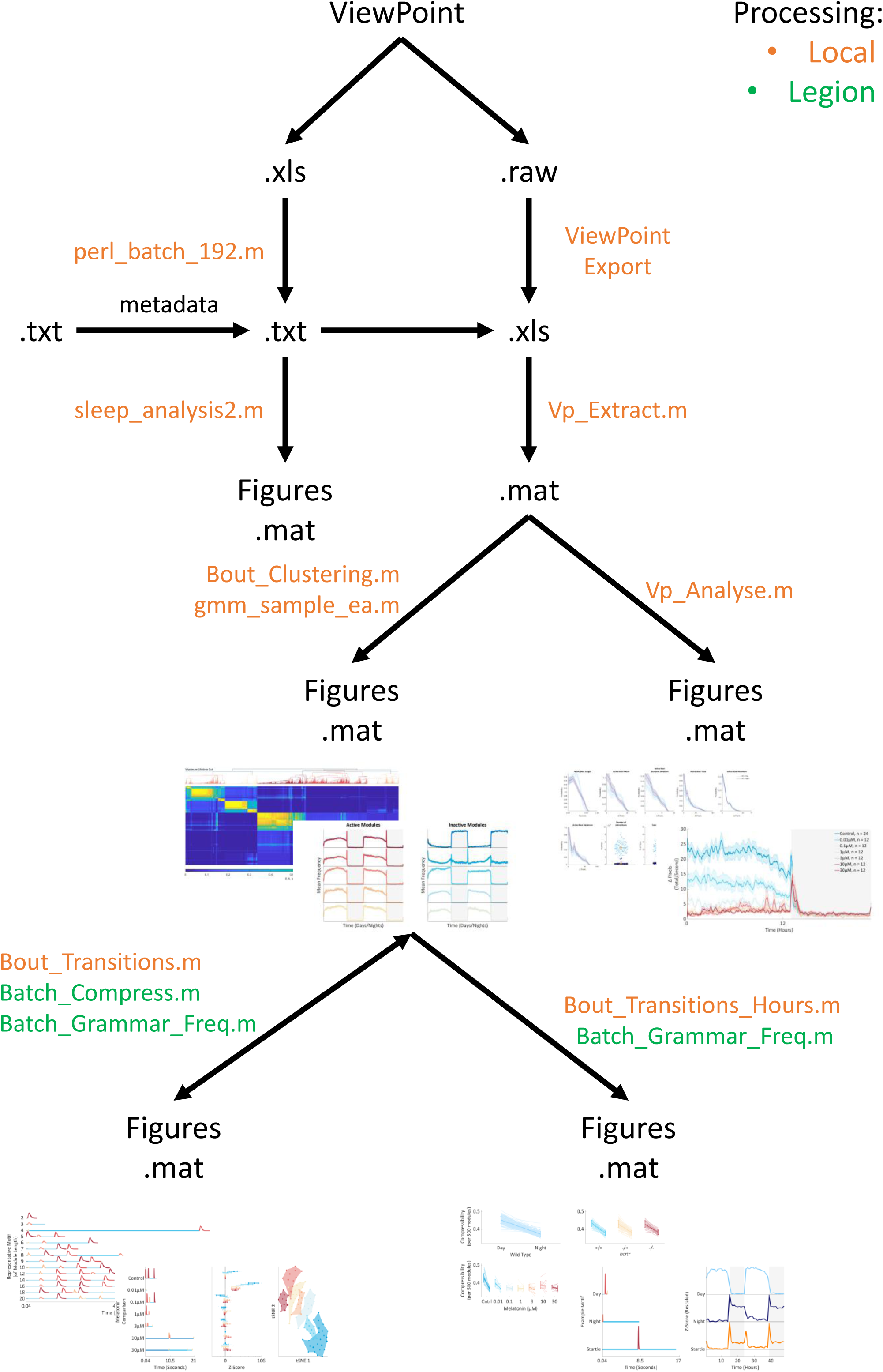
Analysis Framework. Flow diagram depicting the steps of our analysis framework. Data is output from our behavioural set-up (ViewPoint) in the form of a .xls file. perl_batch_192.m organises this data to a .txt format. Experiment metadata (e.g. animal genotypes) is supplied in the form of a .txt file. The 1min bin method uses sleep_analysis2.m to produce figures and statistics from these two .txt files. The 25Hz method exports .raw data from ViewPoint to produce .xls files. Vp_Extract.m reorganises these, using .txt data, to a .mat file which can be input to either Vp_Analyse.m or Bout_Clustering.m. Vp_Analyse.m produces figures and statistics. Bout_Clustering.m uses the clustering function gmm_sample_ea.m to assign data to modules, produce figures and calculate statistics, Bout_Clustering.m’s output can be input to Bout_Transitions.m, which compresses full modular sequences by calling Batch_Compress.m and Batch_Grammar_Freq.m. The motifs identified from this approach can be input to Batch_Transitions_Hours.m which compresses 500 module chunks and uses Batch_Grammar_Freq.m to count motif occurrences per hour. With the exception of the 1min bin method (sleep_analysis2.m), two example figures are shown for each figure producing step. All code can be run locally, though for speed several steps (indicated in green) are best run on a cluster computer.

